# Mapping the human brain’s cortical-subcortical functional network organization

**DOI:** 10.1101/206292

**Authors:** Jie Lisa Ji, Marjolein Spronk, Kaustubh Kulkarni, Grega Repovš, Alan Anticevic, Michael W. Cole

**Affiliations:** Center for Molecular and Behavioral Neuroscience, Rutgers University, Newark, NJ 07102, USA; Department of Psychiatry, Yale University School of Medicine, 300 George Street, New Haven, CT 06511, USA; Department of Psychology, University of Ljubljana, 1000 Ljubljana, Slovenia

**Keywords:** brain networks, brain connectivity, functional MRI, resting-state functional connectivity

## Abstract

- Large-scale functional network map of the entire human brain
- Cortical networks based on multiband fMRI, recently-identified regions
- Subcortical extension of networks covering all subcortical structures
- Multiple quality assessments demonstrate robustness of functional networks
- Network atlas released as public resource, providing framework for future studies

**ABSTRACT:** Understanding complex systems such as the human brain requires characterization of the system’s architecture across multiple levels of organization – from neurons, to local circuits, to brain regions, and ultimately large-scale brain networks. Here we focus on characterizing the human brain’s large-scale network organization, as it provides an overall framework for the organization of all other levels. We developed a highly principled approach to identify cortical network communities at the level of functional systems, calibrating our community detection algorithm using extremely well-established sensory and motor systems as guides. Building on previous network partitions, we replicated and expanded upon well-known and recently identified networks, including several higher-order cognitive networks such as a left-lateralized language network. We expanded these cortical networks to subcortex, revealing 358 highly organized subcortical parcels that take part in forming whole-brain functional networks. Notably, the identified subcortical parcels are similar in number to a recent estimate of the number of cortical parcels (360). This whole-brain network atlas – released as an open resource for the neuroscience community – places all brain structures across both cortex and subcortex into a single large-scale functional framework, with the potential to facilitate a variety of studies investigating large-scale functional networks in health and disease.

## INTRODUCTION

Understanding the highly distributed neural computations that underlie cognitive abilities in humans will require a framework that places neural events in the context of overall brain network organization. Several such frameworks have been introduced (Power et al., 2011; Yeo et al., 2011), based on the idea that the brain exhibits a modular functional architecture (Bullmore and Sporns, 2009). Consistent with this, distant brain regions are strongly functionally interconnected (showing high statistical association between time series), composing distinct functional networks. These functional networks can be detected using resting-state functional connectivity (RSFC) with functional MRI (fMRI), capitalizing on the phenomenon of spontaneous but coherent low-frequency fluctuations of the BOLD (blood-oxygen level dependent) signal. This phenomenon can give insight into the brain’s intrinsic functional network organization that likely underlies a host of computations, including higher-order cognition. This intrinsic organization is thought to be functionally relevant since brain activity patterns during rest and task have high overall correspondence (S. M. Smith et al., 2009). Moreover, task-evoked activity flow (the movement of task-evoked activations between brain regions) can be accurately predicted using RSFC, suggesting that resting-state functional networks provide the pathways over which cognitive task activations flow (Cole et al., 2016a). It is likely also the case that a slower, Hebbian-like learning process allows coactivation patterns to shape RSFC (Harmelech et al., 2013). These hypotheses are additionally supported by findings of strong correspondence between resting-state and task-state functional connectivity: Only subtle changes are observed in brain-wide functional connectivity organization during a wide variety of (functionally distinct) tasks and rest (Cole et al., 2014a; Krienen et al., 2014). These findings suggests that, although smaller task-specific changes in cortical organization occur during tasks, the main functional network architecture is already present during rest.

Based on this idea that intrinsic network organization largely reflects the brain’s functional network organization regardless of state, a number of network partitions have been developed (Doucet et al., 2011; Gordon et al., 2016; Laumann et al., 2015), with two of the most widely-utilized partitions developed by Power et al. (2011) and Yeo et al. (2011). Their widespread impact likely stems from their strong correspondence with well-established primary sensory-motor systems, as well as correspondence with well-replicated co-activation patterns (e.g., frontoparietal co-activations during working memory tasks) in the task fMRI literature (S.M. Smith et al., 2009; Yeo et al., 2015). Both groups used clustering algorithms to identify functional networks based on distributed patterns of high RSFC between brain regions (for Power et al. (2011)) or a grid of cortical surface locations (for Yeo et al. (2011)). Together, they revealed a common brain network organization with bilaterally distributed visual, sensorimotor, default mode, and attention networks. Furthermore, both solutions revealed a task-positive system (the fronto-parietal, dorsal attention and cingulo-opercular networks) and a task-negative system consisting of the default mode network (Fox et al., 2005). These network partitions have proven to be remarkably valuable in elucidating functional brain organization and have provided an initial framework for functional network analyses in a variety of studies, both in health and disease, yielding important new insights in multiple fields of neuroscience (Sporns, 2014).

Yet there are several outstanding issues with existing network partitions, which have limited our understanding of the human brain’s large-scale functional network organization. Most fundamentally, it has been unclear how to define network communities in a principled manner. This difficulty reflects the fact that RSFC values are continuous, with network identification requiring partitioning of these continuous values into labeled clusters. Yeo et al.(2011) were statistically principled in choosing to partition these continuous RSFC values based on the statistical stability of the partition solution at each clustering threshold. Despite this stability, however, there is a major problem with the resulting network partitions: The auditory system is merged with the somatomotor system. The separation of these two systems is a basic property of cortical network organization known for over a century (Fritsch and Hitzig, 1870), and thus serves to question the choice of network partition threshold. In contrast, the Power et al. (2011) partition (and others (Gordon et al., 2016)) has the auditory system separated from the somatomotor system. However, Power et al. (2011) reported nine partitions, with only the most strict cluster threshold yielding a separate auditory system. The nine Power et al. (2011) partitions were subsequently combined into a single partition (Cole et al., 2013; Power and Petersen, 2013) for use in subsequent studies – in a similar manner as a recent surface-based network partition by Gordon et al. (2016). Critically, consensus was achieved across the partitions by hand, raising potential issues with reproducibility. More fundamentally problematic, however, is the possibility that this consensus partition combined networks from distinct levels of organization, by merging network clusters across distinct cluster thresholds. This calls into question the equivalence of the networks in the Power et al. (Power et al., 2011) (and (Gordon et al., 2016)) partition, with potentially problematic implications for studies that utilize this partition (e.g., comparisons of functional systems with functional sub-systems).

In the present study – in light of the primary goal to develop and publicly release a network partition that would be useful to the neuroscience community – we sought to combine the statistically principled approach of Yeo et al. (2011) with the neurobiologically principled approach of Power et al. (2011) (in which the auditory system was separated from the somatomotor system). Specifically, we sought a single-threshold network partition that was statistically stable and that was calibrated to a neurobiologically and functionally meaningful level of organization – the large-scale functional systems level – based on emergence of large-scale sensory-motor systems during network community detection. We chose these systems (visual, auditory, somatomotor) – as opposed to any of a variety of potential networks in association cortex – because (unlike these other networks) the sensory-motor systems are established beyond question. We could thus focus on identification of these networks and be confident that we had identified a fundamental functional systems level of organization even in association cortex, where it remains unclear which networks are fundamental functional systems. Subsequent to identifying our initial network partition, we also used identification of networks reported in the recent RSFC and task activation literatures as supplementary evidence of network partition quality. However, given our principled approach to identifying the functional level of organization, any previously-identified networks that were not identified here were considered likely to be at a distinct level of organization (e.g., the salience network (Seeley et al., 2007) could be a sub-system within a superordinate cingulo-opercular network (Dosenbach et al., 2007)).

As a secondary goal to developing this statistically and neurobiologically principled approach to network detection, we sought to address several methodological limitations of earlier partitions. First, as a starting point, we leveraged a recently-developed surface-based cortical parcellation (Glasser et al., 2016), which combined multiple neuroimaging modalities (i.e., myelin mapping, cortical thickness, task fMRI, and RSFC) to improve confidence in cortical area assignment. Similar to another recent brain region parcellation (Fan et al., 2016), these improvements were driven by convergence across independent MRI-based modalities, each with complementary strengths and weaknesses. Second, we used multiband fMRI data from the Human Connectome Project (Van Essen et al., 2013), allowing for higher spatio-temporal resolution (i.e., simultaneous acquisition of multiple slices at a small voxel size) relative to data used for most previous network partitions (Feinberg et al., 2010; Moeller et al., 2010). This increased the spatial and temporal detail of the RSFC estimates and the resulting network partition. Third, we increased the spatial specificity and anatomical fidelity of the group-level RSFC data by using a high-resolution surface-based analysis in combination with multimodal (functional and anatomical) alignment across subjects (Robinson et al., 2014). Such surface-based methods – especially in combination with multimodal alignment – yield improved cross-subject alignment of cortical geometry (Anticevic et al., 2008; Glasser et al., 2013; Robinson et al., 2014) (relative to volume-based methods). This collectively results in less spatial blurring across sulcal boundaries within an individual and superior cross-areal alignment across individuals (Uğurbil et al., 2013). Notably, several recent cortical network partitions made use of cortical surface data (Gordon et al., 2016; Laumann et al., 2015), demonstrating the utility of this approach for cortical network identification and facilitating the use of surface data in the present study.

Building on these methodological advances, a principled network partition should have a neurobiologically plausible number of networks, including well-known functional systems (Mesulam, 1998; Ryali et al., 2012; Stark et al., 2008) as well as subcortical components (Buckner et al., 2011; Choi et al., 2012). For example, although many known systems were already included in previous network partitions, many of these do not include clear assignment of a language network (e.g. Power et al., 2011). Furthermore, although a few parcellations (e.g. Yeo et al., 2011) have reported the presence of a language network, none have extensively characterized the representation in the subcortex, even though there is ample evidence in the literature for the existence of a distributed language system in humans (Broca, 1861; Hampson et al., 2002; Wernicke, 1874), and perhaps even a homologous network in non-human primates (Mantini et al., 2013). Indeed, this was a major knowledge gap that was partially addressed by the Glasser cortical parcellation – namely identification of putatively novel language-related cortical regions, with clear but heretofore undefined boundaries (Glasser et al., 2016). Therefore, we explicitly tested the hypothesis that a methodologically improved and neurobiologically-plausible network solution should yield a distributed language network based on RSFC graphs. In turn, such a language network should pass the test of mapping onto language-relevant computations based on independent overlap with task-evoked activity during language processing.

We additionally sought to overcome an unresolved technical limitation of previously-developed network partitions whereby there was high uncertainty about the network assignment of the ventral cortical surface. This uncertainty stems from sinus-related MRI dropout (due to magnetic field inhomogeneities) in these regions. The use of multiband fMRI data not only provides a higher signal-to-noise ratio (SNR) due to higher temporal resolution, but also affords less dropout due to higher spatial resolution (Merboldt et al., 2000; Smith et al., 2013). We hypothesized that this would improve network assignments for regions in MRI dropout areas such as orbitofrontal cortex, for which no consensus network assignment currently exists.

Finally, a fundamental knowledge gap in the field is a lack of a unified whole-brain network partition, which includes all of cortex and subcortex. Prior work utilized the Yeo et al. (2011) cortical network assignment to delineate network partitions for the striatum (Choi et al., 2012) and the cerebellum (Buckner et al., 2011), which revealed a shared functional topography between these large anatomical structures (i.e., cortex, striatum, and the cerebellum). However, no study has extended this approach simultaneously across the striatum, cerebellum, thalamus and the brainstem in a common framework. More generally, there is currently no network partition of the entire human brain that capitalizes on the aforementioned cortical mapping advances and that concurrently provides a comprehensive subcortical network mapping. To address this knowledge gap, we derived a network assignment of all subcortical voxels, which we mapped using a connectivity-based clustering analysis building on our cortical network solution (similar to (Buckner et al., 2011; Choi et al., 2012), but including the entire subcortex). The obtained quantitative relationships between subcortical units (voxels) and cortical regions thus help clarify the functional organization of subcortical structures in the context of cortical brain systems.

In summary, the primary purpose of the present effort was to develop and publicly release a brain-wide network partition for the broader neuroscience community. Key innovations beyond prior work include: i) A highly neurobiologically principled definition of cortical network organization, based (in part) on identification of the extremely well-established sensory and motor systems. This criterion was used to calibrate the clustering threshold that identified the brain’s fundamental functional level of organization (i.e., sensory and motor systems). ii) Use of multimodal inter-subject alignment along with surface-based cortical analysis across hundreds of subjects for enhanced precision of the network partition, iii) Use of task and hemispheric lateralization tests to determine reassignment of the previously-labeled “ventral attention” network (Power et al., 2011) as the language network, and iv) Extension of the cortical network assignments to the entire subcortex, providing a whole-brain network solution that expands on prior work that focused on the striatum (Choi et al., 2012) and cerebellum (Buckner et al., 2011). While future improvements of the reported Cole-Anticevic Brain-wide Network Partition version 1.0 (CAB-NP v1.0) are anticipated, this initial solution is shared with the neuroscience community to facilitate studies investigating large-scale functional networks in human health and disease (https://github.com/ColeLab/ColeAnticevicNetPartition).

## METHODS

### Experimental Model and Subject Details

#### Subjects and Dataset

The analyzed dataset consisted of 337 healthy volunteers from the publicly available Washington University – Minnesota (WU-Min) Human Connectome Project (HCP) data (Van Essen et al., 2013). Complete details of all HCP acquisition can be found online (https://www.humanconnectome.org/storage/app/media/documentation/s900/HCP_S900_Release_Reference_Manual.pdf). Further details about the dataset and preprocessing methods can be found in the **Supplementary Materials**.

#### Quantification and Statistical Analysis

The methodological workflows for creating cortical and subcortical network partitions are displayed in **Fig. 2a** and **Fig. 4a**. Specifically, for both partitions, data were first preprocessed via HCP convention, followed by calculation of an average FC matrix (parcel-to-parcel for cortical data or parcel-to-voxel for subcortical data). A cortical partition was then calculated using a clustering algorithm and several pre-determined (hard and soft) criteria, followed by a quantitative evaluation of network solutions. The initial cortical assignment steps are described in detail in forthcoming sections. In turn, this cortical network partition was then used to calculate subcortical network assignment. Here the subcortical voxel was assigned to the cortical network with which is was most highly correlated on average, followed by several quality assurance steps described below.

#### Resting-State Cortical FC Matrices

To sample data at the regional level, we used a recently-developed cortical parcellation (Glasser et al., 2016), which contains 180 symmetric cortical parcels per hemisphere. This parcellation is defined in terms of surface vertices and is thought to be more accurate than prior parcellations due to the consistency of areal borders between data from different modalities and an accurate representation of cortical geometry for each subject via the CIFTI file format (Glasser et al., 2016). Each parcel varied in size and shape based on alignment between functional and anatomical borders across multiple imaging modalities. See Glasser et al. (2016) for details regarding region size and shape. For each subject, BOLD time courses were extracted from the 360 independently identified parcels using Workbench. An average BOLD time course for each parcel was calculated by averaging across all vertices/grayordinates within that region. Subsequently, RSFC between each pair of parcels was calculated for each subject using Pearson correlation. A functional connectivity matrix for N regions is defined as the N×N matrix M, where M(i, j) contains the Pearson correlation of the time courses between region i and region j. In this way, a 360 × 360 RSFC matrix was formed for each subject. Finally, a single group average RSFC matrix was formed by averaging across all subjects in the cohort, and setting the diagonal to zero.

#### Network Detection Using Clustering Algorithm: Louvain Clustering Algorithm

We sought to establish a neurobiologically principled approach to community detection driven by minimal assumptions and excluding qualitative decisions. Our approach was based roughly on (Cole et al., 2014a), which was adapted from methods proposed by Power et al. (Power et al., 2011). We identified three “hard” criteria for what we considered a principled network partition solution, with two additional “soft” criteria.

The hard criteria, as described in the Introduction, included: i) separation of primary sensory-motor cortical networks (visual, auditory, and somatomotor) from all other networks. This criterion is based on unequivocal evidence supporting the existence of these as functionally distinct (and fundamentally functionally meaningful) sensory and motor systems in the human brain. If a network partition is to be neurobiologically-grounded it should pass this standard. Note that previous functional network partitions of the human brain have had difficulty separating auditory cortex from somatomotor cortex (Yeo et al., 2011). Consistent with these prior observations, auditory cortex tended to be merged with somatomotor cortex for most of the tested algorithms and algorithm parameters. Notably, because the auditory network emerged only at a high resolution parameter (after most other potential networks of interest emerged), using any of a wide variety of known large-scale networks (e.g., the DMN or FPN) as additional hard criteria would not have altered the results. Further, the ease of identifying primary sensory-motor networks (due to their extensive characterization from over a century of neuroscientific investigation) suggests that this criterion will be highly replicable in future studies. ii) high stability (similarity of network partitions) across nearby parameters in the network detection algorithm. This criterion served as a heuristic for detecting likely low-noise-influenced partition solutions. iii) High modularity (high within-network connectivity relative to between-network connectivity). This final criterion is implicit in community detection algorithms, which attempt to optimize network partitions for modularity. However, we included this as an additional explicit quantitative criterion to ensure that optimizing for other criteria did not reduce modularity substantially. A putative network solution had to meet the three “hard” criteria to even be considered.

The two “soft” criteria for network partition selection included: i) We optimized the network partition with the constraint that the number of large-scale functional networks should be roughly similar to the number of networks identified in previous functional network solutions using RSFC data (Power et al., 2011; Yeo et al., 2011). These ranged from 7 networks to 17. Importantly, this number of networks is largely consistent with the number of networks typically described in the human fMRI task activation literature, as well as the number of large-scale systems described in the animal neuroscience literature. Put differently, while statistically possible, a network partition with an order of magnitude finer granularity (e.g. >100 sub-networks) would not be considered. ii) We sought a network partition with non-primary networks (other than primary sensory-motor cortical networks that were part of the “hard” criteria our partition, e.g., frontoparietal cognitive control network, default-mode network) qualitatively similar to those that were previously identified using RSFC and fMRI task activations (Power et al., 2011; S. M. Smith et al., 2009; Yeo et al., 2015, 2011). Critically, these two soft criteria had only minimal influence on the finalized partition, since only the hard criteria were used to identify that partition. Instead, these criteria were more important for assessing community detection algorithms, wherein we determined if a given algorithm was providing results (without full parameter optimization) largely consistent with the RSFC, fMRI task activation, and animal neuroscience literatures. Notably, RSFC, fMRI task activation, and animal neuroscience all have weaknesses that are largely non-overlapping (e.g., movement confounds RSFC more than fMRI task activations), such that considering constraints across these sources of evidence strengthens our conclusions.

We started by applying the described criteria across a variety of community detection algorithms. Among the different algorithms explored were OSLOM (Lancichinetti et al., 2011), k-means, hierarchical clustering, SpeakEasy (Gaiteri et al., 2015), InfoMap (Rosvall and Bergstrom, 2008), and the Louvain algorithm (Blondel et al., 2008). Ultimately, the Louvain clustering algorithm method was selected for its ability to easily adjust the resolution of community clustering (i.e., the tendency for smaller communities to be detected), which allowed for optimization of the community clustering based on the “hard” criteria described above. This algorithm was also selected because produced solutions exhibited evidence in support of the “soft” criteria – a number of communities that were broadly similar in number and configuration to what was found in previous RSFC studies and in meta-analyses of fMRI task data. Briefly, the Louvain algorithm works in the following way: First, it searches for small communities by optimizing local modularity. Second, it combines small communities into nodes and builds a new network. Finally, this process is iteratively repeated until modularity changes minimally. Note that we used a modified version of the Louvain algorithm that can accomodate weighted graphs with both positive and negative weights (Rubinov and Sporns, 2011), which allowed us to avoid thresholding the RSFC data used by the algorithm. Ultimately, as with other community detection algorithms, the Louvain algorithm attempts to optimize for the strength of within-community connections relative to the strength of between-community connections (i.e., modularity) (Blondel et al., 2008).

#### Iterative Louvain Clustering and Cluster Consolidation

We started by using a gamma (partition resolution) parameter of 1.0, since this is used as a standard resolution for the Louvain algorithm (Blondel et al., 2008). Initially, this parameter yielded a network partition with the auditory network merged with the somatomotor network, violating one of our “hard” criteria. We therefore initiated a search over gamma values based on the hard criteria described above. As a randomly seeded algorithm dependent on optimization, it is possible that one iteration of Louvain would fail to identify the global (or a near-global) maximum for community modularity. To address this issue, we ran 1000 randomly-initialized iterations of Louvain for each gamma value (range of 1.2 to 1.4 in increments of 0.005), using the Rutgers-Newark supercomputing cluster (Newark Massive Memory Machine). We assessed partition quality by quantifying the stability of the partition solution at each gamma value. Stability estimates were computed as the z-rand partition similarity (Traud et al., 2011) averaged across all iterations for a given gamma value. Thus, if the same parcels were more consistently assigned to the same networks across randomly-initialized iterations for a given gamma value then there would be a higher z-rand score for that gamma value, indicating higher partition stability. The randomly-initialized iteration with the highest average z-rand (i.e., highest mean similarity with all other iterations) was selected as the representative partition solution for that gamma value. Z-rand scores and a calculated modularity score for each generated partition were subsequently examined in the gamma stability analysis described below.

#### Partition Stability Calculation

For each gamma value, z-rand scores and modularity scores across all iterations were averaged to find representative z-rand and modularity values. Next, each mean z-rand score (quantifying partition stability across 1000 iterations) was multiplied by its corresponding modularity score to find a modularity-weighted z-rand score. The gamma value corresponding to the peak of the modularity-weighted z-rand score plot was selected, constrained by the criterion of finding a plausible number of networks including primary sensory/motor networks (see **Fig. 2b**). The partition corresponding to this gamma value was further evaluated for validity and stability across a number of metrics detailed below.

#### Functional Network Validation and Quality Assessment for the Cortical Network Partition

To test the reliability of our network partition (in a distinct manner from the gamma stability analysis), we conducted an independent split-half validation analysis across two randomly selected subsamples of participants. The network detection algorithm was repeated with the same gamma value (1.295) that provided the initial partition solution (based on separation of sensory-motor systems, optimal modularity, and gamma stability), but now with two separate subsets of the data (N=168 and N=169) consisting of demographically matched subjects (see Results section for details on these matched data subsets).

To further quantitatively assess the final cortical network partition and validate the parcel assignments, we used several additional measures. First, a network assignment confidence score was calculated for each region to express the certainty with which that region could be assigned to a particular network (Wang et al., 2015). This confidence score was computed as the difference between the assigned network’s correlation value and the out-of-network correlation values for a region *i*:

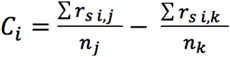

where C_*i*_ is the network assignment confidence score for region *i* (one of 360 brain regions), r_s i,j_ is the Spearman correlation coefficient between the RSFC patterns of region i and region j in the same network, n_j_ is the total number of other regions in regions i’s network, r_s i,k_ is the Spearman correlation coefficient between RSFC patterns of region i and region k outside of regions i’s network, and n_k_ is the total number of regions outside region i’s network. If a region’s RSFC pattern is very similar to that of the other regions in its assigned network, the confidence score will be high, but if it is also similar to other networks, the confidence score will be lower.

Second, in addition to these network assignment confidence scores, we calculated signal-to-noise ratio (SNR) and participation coefficient (Rubinov and Sporns, 2010) (https://sites.google.com/site/bctnet/measures/list), and correlated these measures to assess whether our network assignment results were affected by SNR (i.e. lower functional connectivity in dropout regions) and/or by nodes with extensive between-network RSFC that violate the assumption of network modularity (as assessed by the participation coefficient).

Third, RSFC pattern asymmetry was calculated to see how similar a region’s or network’s RSFC pattern was to that region’s homologue on the other hemisphere. Based on decades of evidence suggesting strong symmetry across hemispheres in RSFC patterns (Biswal et al., 1995; Power et al., 2011; Yeo et al., 2011), we reasoned that high network symmetry (with the likely exception of the language network) was an indicator of network partition quality. In light of the language network being a likely exception, we additionally used this measure to test the hypothesis that there would be especially high (left-lateralized) asymmetry for the language network. For each subject, we correlated each region’s (unilateral) RSFC pattern with that of its homologue (as identified by (Glasser et al., 2016)), and subtracted this value from 1. We subsequently averaged these RSFC pattern asymmetries by network. Finally, scores were averaged across subjects.

Fourth, a measure of inter-subject connectivity variability was used to indicate how similar a region’s functional connectivity pattern is across subjects. To calculate a region’s inter-subject connectivity variability, the rank correlation for each subject’s RSFC pattern for a given region with all other subjects was calculated, resulting in a 337 × 337 (number of subjects X number of subjects) connectivity matrix. Averaging all values in this matrix to generate a mean pairwise similarity score “S” for each region, and subtracting this score from 1, resulted in a region’s inter-subject connectivity variability score (1-S).

Once these quality metrics were calculated, each parcel was assessed for reassignment (i.e. assigning a parcel to a different network than the one resulting from the Louvain clustering algorithm, based on quantitative assessment of its original assignment). We reassigned parcels if the reassignment increased their confidence scores. Reassignment was applied for only three of the 360 cortical regions. Two left hemisphere DMN regions (regions 26 and 75) were re-assigned to LAN, and one left hemisphere LAN region (region 135) was re-assigned to VMM (due to this resulting in higher confidence scores for all three regions). The left hemisphere LAN region (region 135) also failed to replicate on the split-half test, further indicating a poor initial assignment. The reported quality metrics for the partition were recalculated after reassignment of those three regions.

#### Subcortical Network Assignment

Once the cortical network partition was finalized, the subcortical assignment was computed using the cortical partition as a reference. To assign subcortical voxels to networks, FC matrices were first created for each subject containing the correlations between the 360 cortical parcels and 31870 grayordinates covering the entire subcortical CIFTI space. The group FC matrix was then calculated by averaging Fisher’s z-transformed Pearson correlation values across subjects.

Next, the FC of each subcortical grayordinate (i.e. gray-matter vertices or voxels) was averaged across all parcels in each cortical network. In turn, the grayordinate was assigned to the network with the highest mean Fisher’s z-transformed correlation. This approach was chosen to account for the differences in cortical network size, as an unweighted approach would result in a bias towards networks with more cortical parcels.

To account for any signal bleed-over from the adjacent cerebral cortex or partial volume effects in the cerebellum, we removed cerebellar voxels within 2mm of the cortex from the initial network assignment (**Supplemental Fig. S1**). These effects were not prominent in other subcortical structures. We additionally performed cleanup steps of the raw network assignment due to low confidence in making inferences from very small clusters in fMRI data. To achieve cleanup, we removed isolated single-voxel parcels that did not share a network assignment with any adjacent voxels, and parcels of size 2-4 voxels that did not have a counterpart with the same network assignment within a 2mm radius in the contralateral hemisphere. The total number of voxels removed by this process and the map of removed voxels are given in **Supplemental Fig. S2**. We also searched for 5-voxel parcels that would be removed under the same criteria. The difference achieved with the 5-voxel criteria was minimal. An additional 5 voxels, or 0.016% of the total subcortex, was flagged in the 5-voxel version (**Supplemental Fig. S2**), suggesting that the 4-voxel version was already fairly stable. To provide a complete functional atlas of the entire subcortical space, we used nearest-neighbour interpolation to reassign the voxels removed from network assignment in the previous steps. Lastly, parcels which shared a corner and had a continuous contralateral counterpart were combined.

#### Subcortical Network Assignment of Data with and without Global Signal Regression

Given the well-known concerns surrounding artifact removal from BOLD data, we also computed two a solutions of the subcortical network assignment using BOLD signal with and without global signal regression (GSR) performed on top of the canonical HCP-style de-noising (i.e. minimal preprocessing + FIX ICA, (Glasser et al., 2016)). Specifically, GSR was performed by regressing the global mean gray matter signal from each resting-state BOLD time series for each subject. We then repeated the network assignment for subcortical voxels using the same procedure as described above (“Subcortical Network Assignment”). The similarity between the two versions was quantified by calculating the proportion of each network in the original partition that was stably replicated with the GSR version (see **Supplemental Fig. S4**). Chance overlap was calculated using the hypergeometric test, as described below above (“<u>Evaluating the Subcortical Network Assignment</u>”).

#### Evaluating the Subcortical Network Assignment

The stability of the subcortical network assignment was tested using a split-half validation analysis, similar to the procedure performed for the cortical network partition. The same network assignment steps described above were performed independently for two separate samples (N=168 and N=169) consisting of matched subjects (**Fig. 4a-b**). To quantitatively compare the discovery and replication solutions, the proportion of voxels which were assigned to the same network in both solutions was computed. This was done before and after the described cleanup steps were performed (**Fig. 4e-d**). The proportion of voxels expected to overlap by chance in both solutions was calculated for each network, by using the hypergeometric test for proportions given the total number of voxels in the network and the total number of all subcortical voxels. 95% confidence intervals for chance were calculated with the Clopper-Pearson method (Clopper and Pearson, 1934).

Additionally, the asymmetry of the subcortical partition was evaluated. Asymmetry was also computed voxelwise because the network assignment for the subcortex was computed on per-voxel level, rather than per-parcel level (as with the cortex). A homologous pair of subcortical voxels was defined such that they had to be equidistant along the x-axis relative to the midline (y-axis). Observed symmetry was computed as the proportion of voxels in each subcortical network for which the homologous voxel was assigned to a different network (**Fig. 5e**). Chance asymmetry was calculated as the proportion of voxels in each network that would be expected to overlap between left and right hemispheres if the voxels were randomly assigned, given the number of voxels in the network and the total number of voxels in the subcortex. Additionally, the proportion of each subcortical network in the left and right hemispheres was computed (**Fig. 5f**). Because anatomical connections to and from the cerebellum cross the midline at the level of the pons (van Baarsen et al., 2016) and functional representation (e.g. of somatotopic maps) is mirrored relative to the rest of the brain, the left and right cerebellar hemispheres were exchanged in this analysis.

#### Task Activation fMRI Analyses

We evaluated the correspondence of the identified networks using task fMRI data from a language processing task and a motor task in the same sample of subjects. Briefly, the language processing task consisted of two runs, each with 4 blocks of a ‘LANGUAGE’ processing task, which consisted of three components: (i) Auditory sentence presentation with detection of semantic, syntactic and pragmatic violations; (ii) auditory story presentation with comprehension questions; (iii) Math problems that involved sets of arithmetic problems and response periods. Both the ‘Story’ and ‘Math’ trials of the LANGUAGE task were presented auditorily and participants chose one of two answers by pushing a button. Further details concerning the LANGUAGE task have been previously described in full by Barch and colleagues (Barch et al., 2013; Binder et al., 2011). Notably, Glasser and colleagues (2016) demonstrated that Area 55b, defined through multi-modal parcellation, was robustly activated in the ‘Story versus Baseline’ task contrast from the HCP’s ‘LANGUAGE’ task. Here we leveraged that contrast to validate the language system. Specifically, task-evoked signal for the LANGUAGE task was computed by fitting a general linear model (GLM) to preprocessed BOLD time series data. Two predictors were included in the model, for the ‘Story’ and ‘Math’ blocks respectively. Each block was approximately 30s in length and the sustained activity across each block was modeled (using the Boynton hemodynamic response function (Boynton et al., 1996)). In turn, three unique contrasts were computed for the LANGUAGE task: i) Story versus Baseline, ii) Math versus Baseline, and iii) Story versus Math. Here we focused on the ‘Story versus Baseline’ contrast, as reported by (Glasser et al., 2016). The motor task used in the HCP consisted of two runs, based on the task protocol used by (Power et al., 2011; Yeo et al., 2011) and (Buckner et al., 2011; Choi et al., 2012). Each run was composed of 13 blocks: 3 fixation blocks, 2 blocks of tongue movements, 4 of hand movements (2 right and 2 left), and 4 of foot movements (2 right and 2 left), with each block lasting 12 seconds (10 movements). Participants were given a 3 second visual cue at the start of each block to signal which body part to move. Here, we computed the foot and hand versus tongue contrasts to suppress visual and attentional effects in the task-related activation. As reported in (Buckner et al., 2011; Choi et al., 2012), while the movement versus fixation contrasts produced similar activation maps, the comparison of two movements removed nonspecific responses in both cortex and subcortex.

### Data and Software Availability

Data, software, and the network partition are available here: https://github.com/ColeLab/ColeAnticevicNetPartition and https://doi.org/10.5281/zenodo.1455791.

## RESULTS

### Cortical Network Partition

The overarching objective of this study was to identify a brain-wide large-scale functional network organization based on clusters of multimodally-defined cortical regions – the likely next-lowest level of organization from the large-scale functional network level (Felleman and Van Essen, 1991; Glasser et al., 2016; Van Essen and Glasser, 2014). Since higher levels of organization emerge from the units at lower levels, increased accuracy in mapping brain regions (lower level) may yield more accurate large-scale brain networks (higher level). We therefore quantified functional networks based on regions recently identified via convergence across multiple modalities via both functional and structural criteria (Glasser et al., 2016), increasing confidence of their spatial precision. This yielded a cortical network organization (**Fig. 1a and Fig. 1c**) largely consistent with known and recently-identified functional networks, along with several previously-unidentified but highly robust networks.

**Figure 1.**
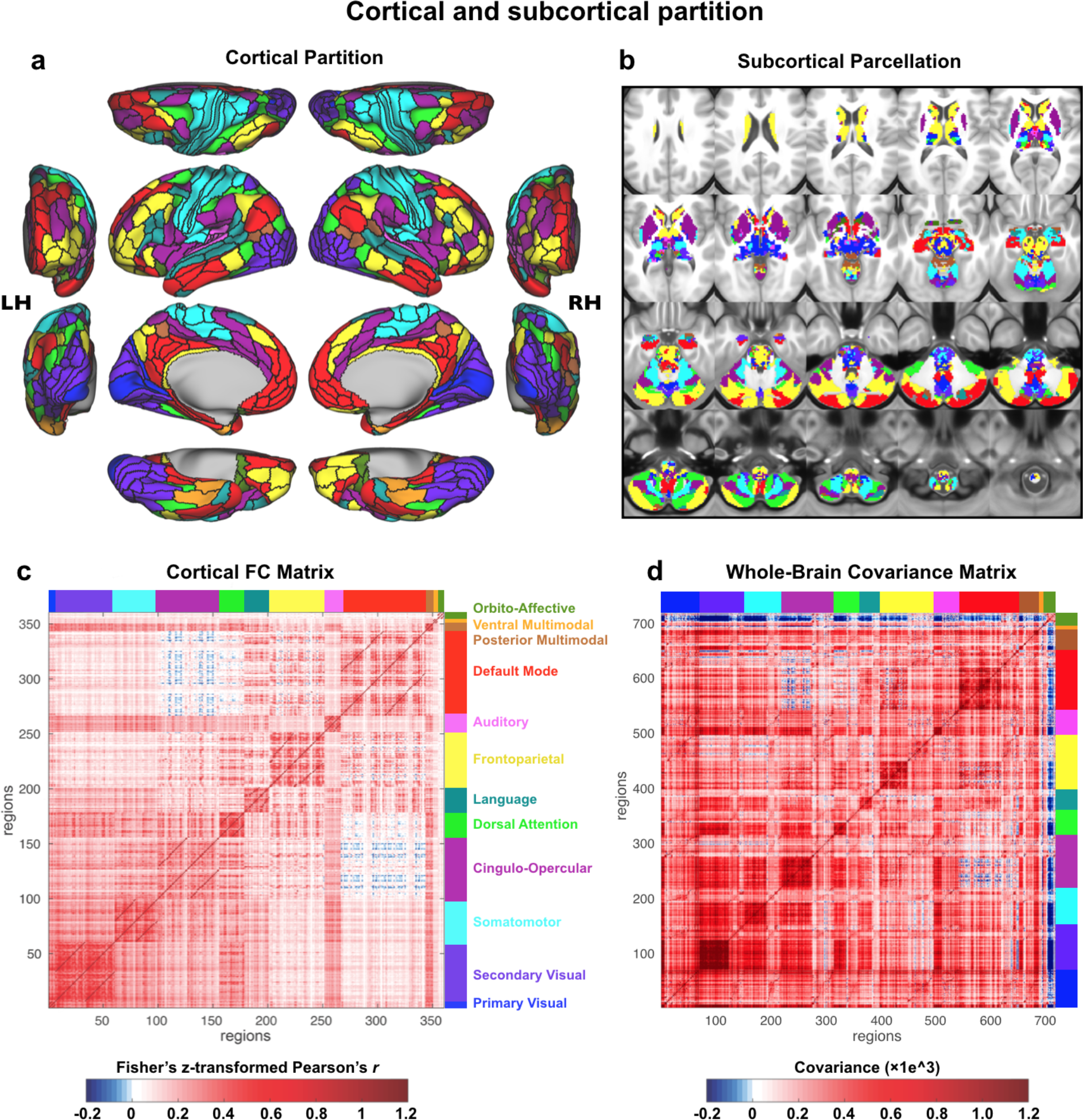
Cortical-subcortical network partition. **A**) The cortical network partition, as calculated with cortical surface resting-state fMRI data using graph community detection. We focused on identifying the network level of organization based on interactions among the next-lowest level of organization – functional regions. Network detection was calibrated based on identification of the well-established primary sensory-motor cortical systems (visual, somatomotor, auditory). Identifying clusters of multimodally-defined cortical regions replicated many known and revealed several novel large-scale networks. **B**) The network partition identified in cortex was extended to all subcortical gray matter voxels. Briefly, each voxel was assigned to the cortical network with the strongest average resting-state functional connectivity (FC) with that voxel. **C**) The region-with-region FC matrix within cortex, sorted by network assignment. The block-like structure along the diagonal provides a visualization of the greater FC strength within (relative to between) each network. The darker off-diagonal lines reflect stronger cross-hemisphere FC within networks (since left hemisphere regions are listed first within each network). **D**) The parcel-to-parcel FC (covariance) matrix, including both cortical and subcortical parcels. Covariance is a non-normalized version of Pearson correlation, used here to account for higher standard deviation of time series in subcortical parcels. We previously validated covariance as a valid alternative to Pearson correlation for FC estimation (Cole et al., 2016b).

Briefly, we used graph community detection to identify clusters of highly interconnected cortical regions based on RSFC (**Fig. 2a**; see **Methods** for details). We used a standard community detection algorithm that identifies communities by optimizing for modularity (high within-network and low between-network connectivity strength) (Blondel et al., 2008). Several principles were used to calibrate the definition of network communities as we searched over different “resolution” (gamma) parameters in the community detection algorithm: i) We required that primary sensory-motor cortical regions (visual, auditory, somatomotor) – which have been known for over a century to be functionally distinct neural systems (Fritsch and Hitzig, 1870) – would be identified as separate functional networks. Such separation was clear at the default “resolution” setting of the community detection algorithm (gamma=1) for separation of visual and somatomotor networks, but the auditory network was merged with the somatomotor network. We therefore increased the community resolution parameter until auditory and somatomotor networks separated. ii) We optimized for stability (similarity of network partitions across neighboring parameter settings) and iii) we optimized for modularity (high within-network and low between-network connectivity strength) (**Fig. 2b**; **Supplemental Fig. S5**). Note that we had a “soft” requirement (see Methods) that the major networks identified in the fMRI task activation and RSFC literatures (DMN, FPN, DAN, and CON) would be present, but these networks emerged with the above criteria (no additional steps necessary). This approach revealed 12 networks consisting of well-known sensory-motor networks, previously-identified cognitive networks, and several novel networks.

Well-known networks included primary visual (VIS1), secondary visual (VIS2), auditory (AUD), and somatomotor (SMN) networks. Previously-identified cognitive networks – networks identified in the last few decades – included the cingulo-opercular (CON), default-mode (DMN), dorsal attention (DAN), and frontoparietal cognitive control (FPN) networks. Two primary functional network atlases were used to identify these previously-identified networks: Power et al. (2011) (which was updated by (Cole et al., 2013)) and Yeo et al. (2011). Novel networks included the posterior multimodal (PMM), ventral multimodal (VMM), and orbito-affective (ORA) networks. We include additional analyses below to better establish the robustness of these networks, given that they have not (to our knowledge) been previously described. Notably, we also identified a language network (LAN), which has been known for over a century (Broca, 1861; Wernicke, 1874), yet has been missing from most previous atlases of large-scale functional networks (e.g. Power et al., 2011) and never extensively characterized subcortically. We include additional analyses below to establish that this network is involved in language functions and is likely equivalent to the previously-characterized left-lateralized language network consisting of Broca’s area and Wernicke’s area (among other language-related regions).

**Figure 2.**
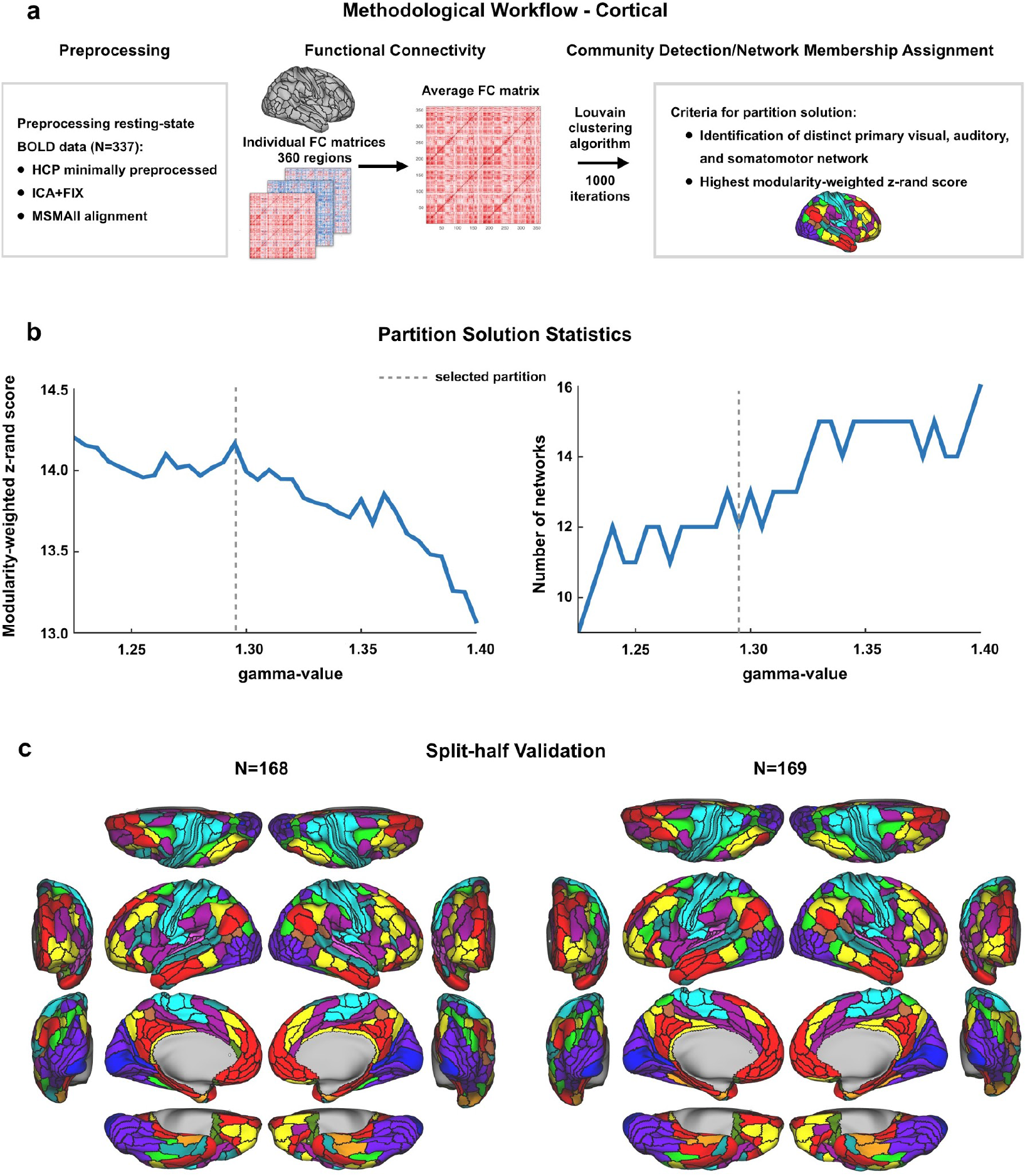
Cortical partition solution workflow and statistics. **A)** Schematic workflow used to create cortical partition. Data were preprocessed for 337 subjects, functional connectivity was calculated between all regions for each subject, and an FC matrix was constructed for each participant. After averaging across subjects, the Louvain clustering algorithm was run with 1000 iterations to detect communities of networks for a range of gamma-values. The final cortical partition was a result of two criteria; a plausible number of networks that included primary sensory/motor networks had to be present, and the most stable and modular partition solution was chosen. **B)** Plots presenting the modularity-weighted z-rand scores and number of networks in the partition for each gamma-value. The dashed line indicates at which gamma-value the community detection gave the most stable and (neurobiologically) plausible results. **C)** Split-half validation results for the cortical partition. The original dataset was split in two smaller sets consisting of matched subjects’ data and the Louvain clustering algorithm was run with the same parameters as for the original partition. The two resulting network partitions were both highly similar to each other (92.5% overlap in network assignments) and highly similar to the original one presented in **Fig. 1a**, indicating that our partition is reliable. See main text for more details.

### Assessing Quality of Cortical Network Partition

We used a split-half analysis to estimate the reliability of the cortical network partition (**Fig. 2c**). The identical algorithm (with identical parameters) was applied to a pseudo-random set of 168 subjects (selected from the total set of 337 subjects), and then independently to the remaining 169 subjects. The split-half sets were matched on a variety of demographics in order to reduce the chance that observed differences were driven by group differences of potential interest (e.g. age or gender). The 168 subjects were selected by first creating a random list of subjects then exchanging subjects between the groups such the 168 subjects were matched, at the group level, with the remaining 169 subjects on the following demographics: age, gender, handedness, and education. This analysis revealed a highly similar network partition across the the two independent matched samples (**Fig. 2c**): adjusted z-rand (Traud et al., 2011) of z=190.2 (*p*<0.00001) and 92.5% of regions with identical network assignments. These results demonstrate high reliability of the main cortical network partition.

To further quantitatively evaluate the cortical partition, we calculated a network assignment confidence score (**Fig. 3a & c**), inter-subject connectivity variability (**Fig. 3b**), network-level split-half overlap (**Fig. 3d**), and network RSFC pattern asymmetry (**Fig. 3e** **&** **Fig. 8d**) for each parcel and network. As shown in **Fig. 3a & c**, most networks exhibited broadly similar confidence, with a mean score of 0.36 (SD=0.08), indicating higher RSFC pattern correlation between a region and its assigned network than with other networks. Only ORA had a substantially lower confidence score (mean=0.19, SD=0.1), possibly as a result of lower SNR in regions assigned to the ORA network (mean SNR=152 with range 143-194) compared to other networks (mean SNR all networks=228, range 79-371). Note that it is possible for confidence scores to be negative, which would indicate more confidence in an alternative assignment than the one provided by the network partition. Thus, the positive confidence values across all networks indicates accuracy of the network assignments.

A fundamental assumption of network partition analyses is that the brain’s functional network architecture is highly modular (i.e., that it has minimal between-network connectivity). We reasoned that confidence scores – which closely reflect this modularity assumption – would be lower as a function of how much this assumption is violated. Based on this, we hypothesized that low confidence could reflect three potential sources, each driving real or apparent between-network connectivity: low SNR, high participation (Guimerà et al., 2005), or high intersubject variability. First, we reasoned that low SNR would result in additional (though likely weak) random connections, reducing the apparent clustering/modularity of connections and therefore decreasing confidence scores. Second, we reasoned that the modularity assumption would be violated by between-network connector hubs – which would have high partition coefficients (Guimerà et al., 2005; Power et al., 2013, 2011) – since such nodes do not fit neatly into a single network partition. Finally, we reasoned that high inter-subject variability could result in apparent low modularity via averaging connectivity of distinct brain regions (with connectivity to potentially distinct networks), lowering confidence scores. Our primary strategy for testing these hypotheses was to assess the relationship between these three factors and confidence scores.

**Figure 3.**
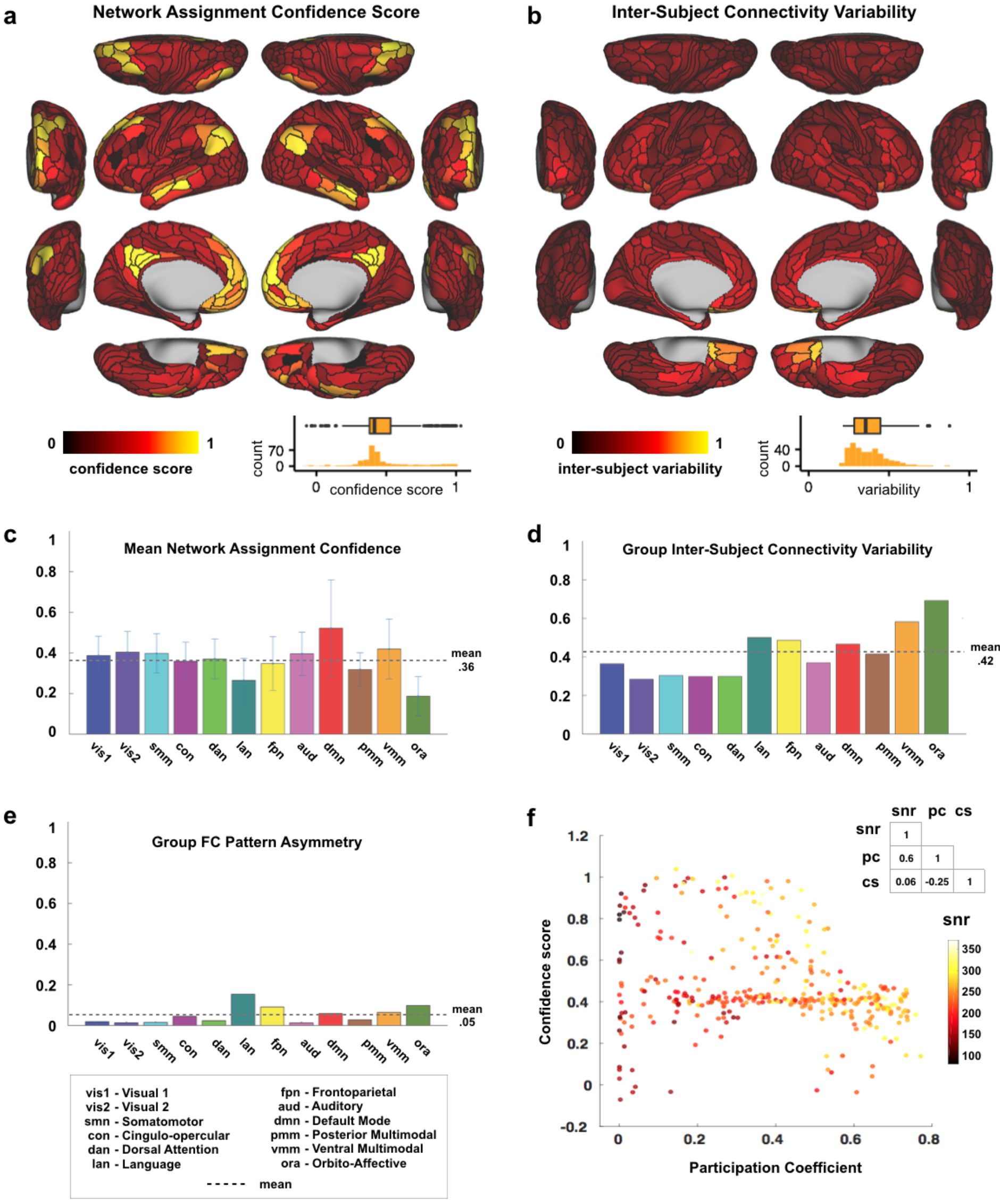
Quantitative assessment of cortical network partition. **A)** Cortical map with Network Assignment Confidence scores, reflecting a region’s fc pattern similarity (calculated using Spearman’s rank correlation) to its assigned network divided by similarity to all other networks. These scores are used as a measure of certainty that the network to which a parcel was assigned is the correct one. The mostly homogeneous map indicates similar confidence across regions. Inset shows the distribution of confidence scores across the 360 cortical regions. **B)** Cortical map displaying Inter-Subject Connectivity Variability, a measure comparing the connectivity patterns for each region across subjects. Similar to panel A, most cortical regions appear to have highly similar values. Inset shows the distribution of intersubject variability across the 360 cortical regions. **C)** Network averages of the parcel-level network assignment confidence scores (in panel A) are displayed. Error bars indicate standard deviations. Highest confidence scores were found in DMN and lowest in the new orbito-affective network (but note the lower SNR in this area). **D)** Split-half replication assignment overlap by network. This quantifies the amount of overlap between the split-halves in **Fig. 2c**. **E)** Group FC Pattern Asymmetry, reflecting similarity between a region’s (unilateral) functional connectivity pattern and that of its supposed homologue region on the opposite hemisphere. Note the relatively high asymmetry for the language network (LAN) resulting from the left-lateralized language parcels in our partition. **F)** Scatterplot showing the relationship between Network Assignment Confidence score, Participation Coefficient and SNR for each parcel. See main text for the logic behind this analysis. The non-significant correlation between Confidence and SNR indicates that Confidence scores were not substantially affected by SNR. However, a negative correlation between Confidence and Participation Coefficient could indicate that lower confidence regions partly consist of connector hubs that are shared between multiple networks (violating modularity).

We evaluated these possibilities, as shown in **Fig. 3f**. SNR and confidence scores were not significantly correlated (r=0.06, *p*=.29), which is inconsistent with the possibility that low SNR strongly affected confidence scores in these data. Regions with higher participation coefficients, a measure indicating how distributed a node’s edges are across networks (potentially violating the assumption of a modular network organization), exhibited lower confidence scores (r=−0.25, *p*<0.00001). This suggests that low confidence might be explained by connector hubs (regions connecting to multiple networks). Note that participation coefficient was calculated on the single-subject level, ruling out the possibility that high inter-subject variability drove the participation results.

Together, these results suggest network assignment quality was primarily influenced by high participation coefficient (strong RSFC with multiple networks) rather than low SNR. This suggests that the human brain violates modularity to some extent, reducing assignment confidence because some regions are connected to multiple networks. Note, however, that connectivity of single regions with multiple networks is not entirely surprising since the brain must somehow integrate functionality between networks, which requires variable inter-network connectivity. These results suggest the degree of multi-network connectivity may be small overall, however, since participation accounts for only 6.25% of the linear variance in confidence scores (participation-confidence r=−0.25; r^2^=0.0625).

We also expected that low confidence could be driven by high inter-subject connectivity variability, which we calculated as the mean dissimilarity of a given region’s cortex-wide RSFC pattern across subject. This could have been driven by the kinds of subject-to-subject variability in RSFC patterns shown in several recent studies (Braga and Buckner, 2017; Gordon et al., 2017). Inconsistent with this being a major factor, we found a relatively homogeneous level of inter-subject variability across the Glasser parcels (**Fig. 3b**), with a mean variability score of 0.42 (SD=0.13) for the networks. Overall, most networks exhibited low inter-subject connectivity variability, relative to a maximum value of 1.0 (in which every subject’s connectivity pattern would differ completely from every other). One network with higher inter-subject variability was the VMM network (mean=0.59, SD of 4 VMM regions=0.006), but this network’s high confidence score suggests its networks assignment are nonetheless accurate overall. The ORA network also showed numerically higher inter-subject connectivity variability between subjects compared to the other networks (mean=0.7, SD of 6 ORA regions=0.15), in concordance with this network’s lower confidence score. Note that rather than true inter-subject variability this may have been driven by somewhat lower SNR (i.e., greater measurement noise) in ORA regions (Spearman correlation between ORA inter-subject variability and SNR: r=−0.25, *p*<0.00001), likely due to MRI signal dropout from nearby sinuses. These results suggest some details are lost by using group-level RSFC (rather than individual-level RSFC) to identify networks, but that most network assignments are likely accurate.

We additionally assessed partition quality by quantifying inter-hemispheric asymmetry, under the assumption that most networks would be highly symmetric across the hemispheres. This assumption is based on the well-established observation that most cortical regions have high RSFC with their homologue in the other hemisphere, despite overall higher within-hemisphere RSFC and some variability in inter-hemispheric symmetry (Stark et al., 2008). This metric served as a “proxy” test of reliability, since we did not constrain the network partition to be symmetric. Asymmetry scores were calculated as the dissimilarity of cortex-wide RSFC patterns across hemispheric homologues (see **Methods**). Asymmetry results in **Fig. 3e**show that for most regions/networks RSFC patterns were very similar to a region’s/network’s homologue on the contralateral hemisphere (network mean=0.05, SD=0.04; all far below complete asymmetry of 1.0). An exception to this, which was expected based on the language neuroscience literature, was the LAN network having the highest cortical asymmetry score. This reflects the left lateralization of this network (see also **Fig. 8** and additional analyses for LAN below), with 14 LAN regions on the left hemisphere vs. 9 regions assigned to LAN by the Louvain algorithm on the right hemisphere. Overall, this result further demonstrates the quality of the network partition, given that all networks showed substantial inter-hemispheric symmetry with the expected exception of the LAN network.

### Subcortical Extension of the Cortical Network Partition

Previous functional atlases of the human brain have focused primarily on cortical network assignments. However, it is well established that vital neural computations are also implemented by subcortical regions. Furthermore, many of the subcortical nuclei form functional loops, via the thalamus, with cortical territories. Thus, we expanded our network mapping to subcortical structures to develop a comprehensive whole-brain functional network atlas. We built on recent efforts to extend cortical network definitions into cerebellum (Buckner et al., 2011) and striatum (Choi et al., 2012), but extended our network assignment to all subcortical structures, additionally including: thalamus, hypothalamus, amygdala, hippocampus, brainstem, and all of basal ganglia, in addition to all other subcortical nuclei (**Fig. 1b**). Together with the cortical partition this yielded a whole-brain solution for large-scale functional networks whose raw covariance matrix we present in **Fig. 1d** and **Supplementary Fig. S6-S7**.

Briefly, we assigned each voxel to the network with which it shared the highest mean connectivity (using Pearson correlation) across cortical parcels. We then implemented a number of quality control cleanup steps to eliminate small parcels that may be noise-driven, or that may have been driven by partial volume effects near the edge of cerebellum (**Fig. 4a**; see **Methods** and **Supplementary Fig. S8** for details). Parcels were also constrained to anatomical boundaries between major subcortical structures, as defined by Freesurfer, to conform to the gross anatomy of the subcortex. We computed a subcortical network solution using both resting-state fMRI data with and without GSR (hereafter referred to as wGSR and woGSR, respectively), due to concerns that low SNR in the subcortex may lead to extensive assignment of voxels to the visual networks (see **Methods**). The wGSR subcortical parcellation produced a largely symmetric solution with 358 parcels, presented in **Fig. 4**. This solution was highly replicable across split-half samples, both qualitatively (**Fig. 4b-c**) and quantitatively (**Fig. 4d-e** see **Methods**). The proportion of voxels that were assigned to the same network in both Discovery (N=168) and Replication (N=169) samples was highly significantly above chance for all networks (**Fig. 4d**).

After quality control cleanup steps were performed for each of these split-half solutions, the proportion of replicated voxels increased for all networks (with the exception of VMM, **Fig. 4e**). Critically, we found that all 12 cortical networks, including higher-order associative networks (such as the FPN and CON), were represented in the subcortex with predominantly symmetrical and robustly replicable assignments.

The woGSR parcellation also resulted in a highly symmetric and replicable solution (**Supplemental Fig. S3** and **Supplemental Fig. S4**), despite the possibility of more noise being present due to global signal artifact. Voxelwise network assignment and cleanup steps were performed identically to the wGSR version, as described above. The woGSR parcellation produced 288 distinct subcortical parcels after all cleanup steps, and showed more extensive assignment of the visual networks. The number of voxels with stable assignments was significantly above chance for all networks given the total number of voxels in the subcortex.

**Figure 4.**
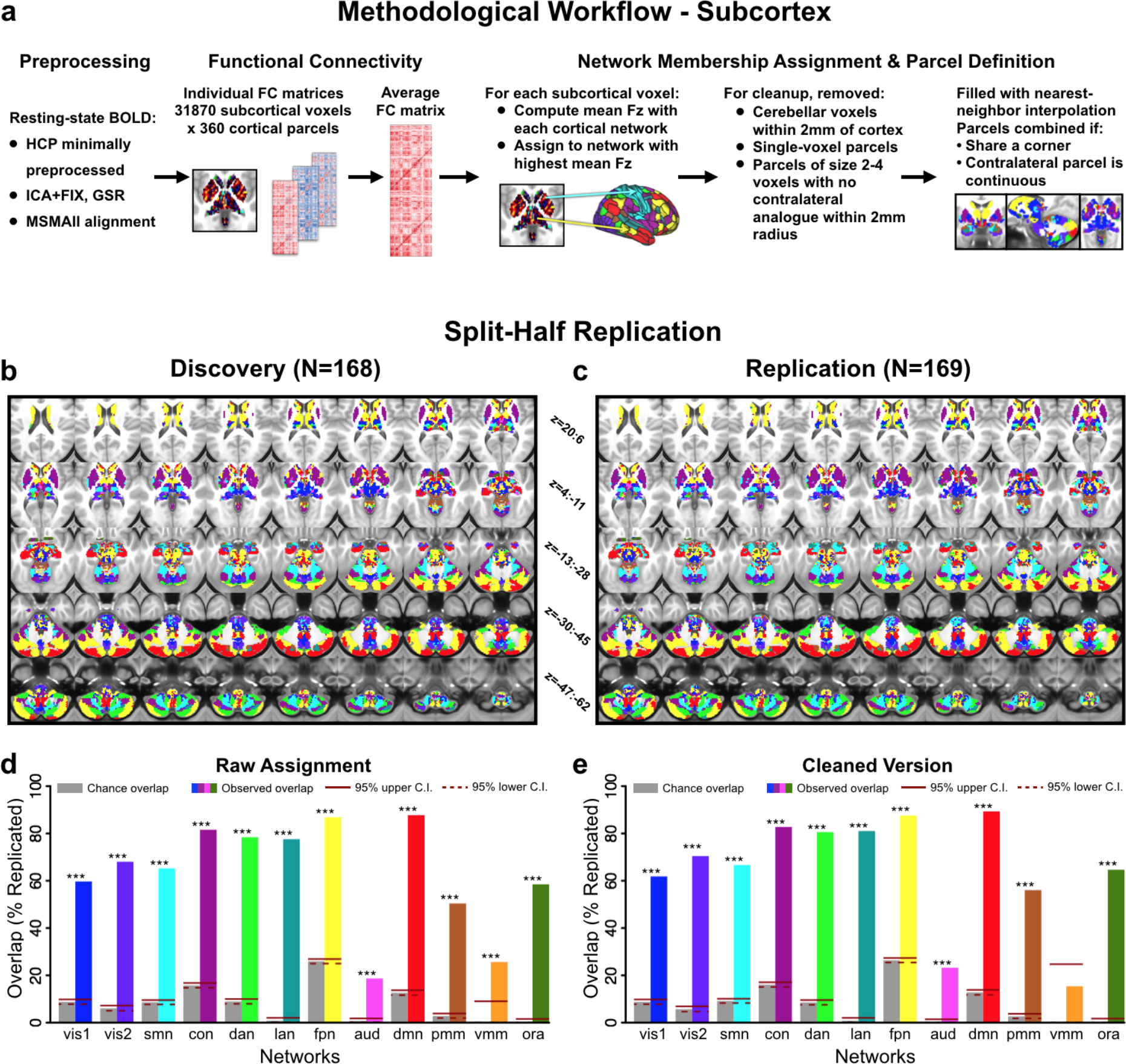
Subcortical partition solution workflow and statistics. **A)** Schematic workflow used to create subcortical partition. **B & C)** Split-half replication of the subcortical partition. The subcortical network assignment procedure was performed independently on two smaller sets of subjects matched for demographic variables. **D)** Proportion of voxels in each network with replicated assignments, before any cleanup steps. Gray bars show proportion of voxels expected to replicate by chance given the size of each network. Solid and dashed red lines indicate upper and lower 95% confidence interval for chance, respectively. **E)** Proportion of voxels in each network with replicated assignments, after cleanup steps were performed (see **Methods**). The proportion of voxels with identical network assignments in both Discovery and Replication samples was significantly above chance for 11 out of the 12 networks (*p*<0.05), suggesting that the subcortical solution is highly replicable.

To additionally verify the subcortical network partition and evaluate the wGSR versus woGSR versions, we compared the network assignments to Motor task activation maps computed in the same sample of subjects. This task has previously been used for localizing motor regions in both the cortex and subcortex (Barch et al., 2013; Buckner et al., 2011; Cole et al., 2016a; Yeo et al., 2011) within the boundaries of the SMN. In the subcortex, while both the wGSR version (**Fig. 5b**) and the woGSR version (**Fig. 5c**) overlap with task-activated regions, the congruence of motor task activation with the wGSR version is higher both qualitatively and quantitatively (**Fig. 5d**). This effect was notable even with single task-related contrasts. As an exemplar, **Fig. 5e** highlights the task activation in the cortex of the left foot (LF) versus Cue contrast, which falls cleanly within the contralateral (right) SMN. In the subcortex, the wGSR version of the parcellation clearly delineates the task-activated area in the contralateral thalamus (**Fig. 5f**) and the ipsilateral (left) cerebellum (**Fig. 5h**). While the SMN in the woGSR version of the parcellation largely occupies the same areas, the overlap with task-activated regions is noticeably less clean (**Fig. 5g**, **Fig. 5i**). Due to this, we present results using the wGSR version of the subcortical parcellation for our remaining analyses, although the woGSR parcellation is also available as part of our public release. The high degree of convergence between our derived whole-brain networks and task-related activation patterns is a strong indication that these networks (even in the subcortex, where FC values were lower) are functionally relevant.

**Figure 5.**
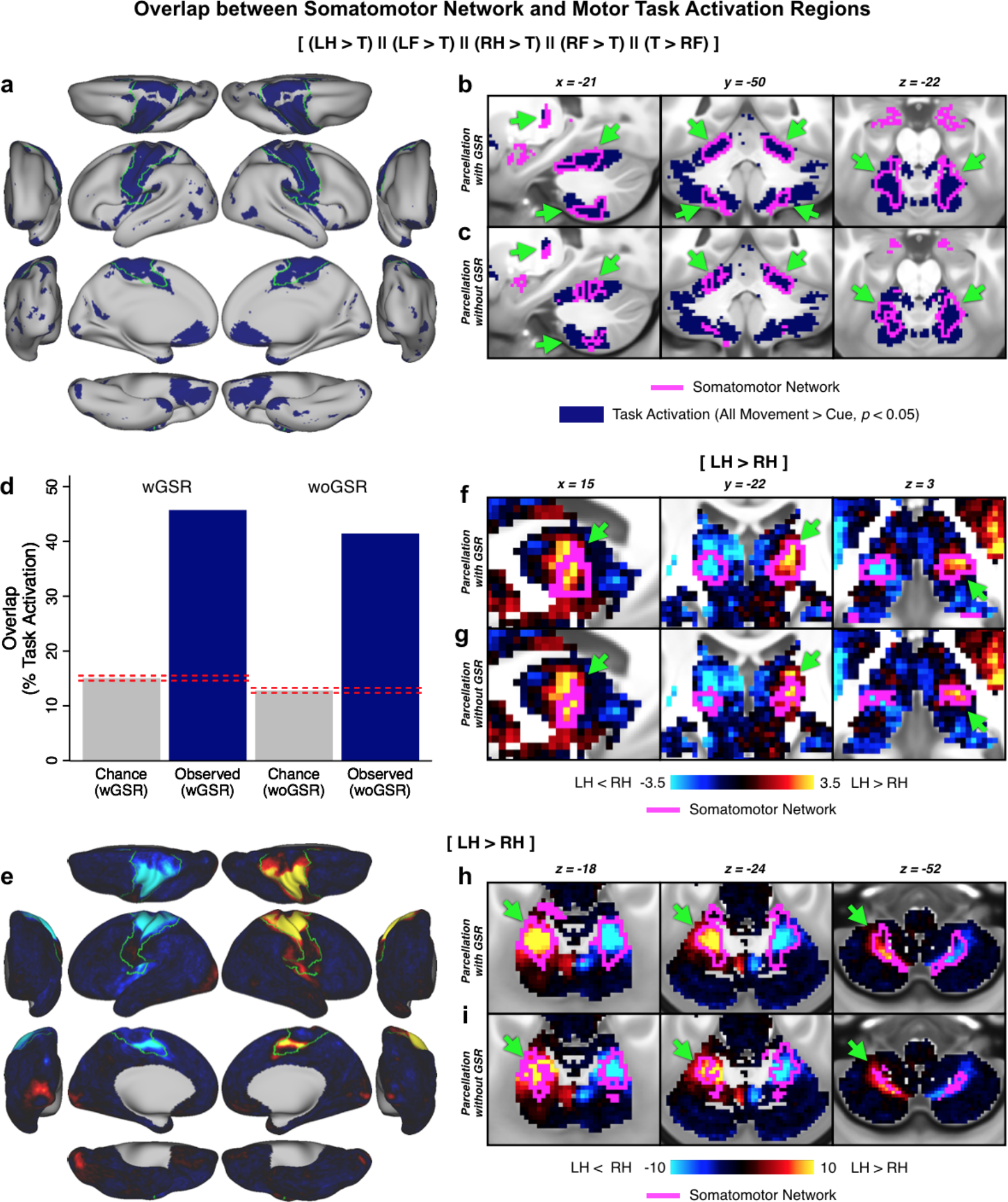
Convergence of cortical and subcortical network partition and motor task activation. The motor network is shown as evidence for valid extension of cortical functional networks to subcortical regions. **A)** Combined motor task responses for comparisons between two movements [(left foot > tongue), (left hand > tongue), (right hand > tongue), (right foot > tongue), and (tongue > right foot)] in the cortex, with the SMN outlined in green. **B)** Combined motor task responses in the subcortex, with the SMN from the wGSR subcortical parcellation outlined in fuchsia. Arrows highlight regions of convergence between task activation and SMN. **C)** Same data as **B** but with the woGSR SMN. **D)** Comparison of overlap between subcortical task activation and subcortical SMN from the wGSR and woGSR partitions. Dashed lines indicate 95% confidence interval for chance. Because the degree of convergence is higher for the wGSR version, we use this for all subsequent subcortical analyses presented in this study. **E)** Map of the left foot (LF) > tongue (T) contrast in the cortex, with the SMN outlined in green. **F)** Map of the left hand (LH) > right hand (RH) contrast in the thalamus, with the SMN from the wGSR subcortical parcellation and **G)** the SMN from the woGSR subcortical parcellation outlined in fuchsia. **H)** Map of the LH > RH contrast in the cerebellum, with the SMN from the wGSR subcortical parcellation and **I)** the SMN from the woGSR subcortical parcellation outlined in fuchsia. Note the ipsilateral representation of the hand movements in the cerebellum and the higher convergence of the wGSR parcels relative to the woGSR parcels with task activation.

Importantly, prior subcortical network assignment attempts did not incorporate the thalamus and the brain stem in their reported solutions. As noted, thalamic subnuclei are well-known to form functional circuits with cortical networks (Barbas, 2000; Zhang et al., 2008) and have been shown to exhibit robust patterns of diffusion MRI-derived probabilistic tractography with cortical territories (Behrens et al., 2003). Therefore, it was vital to demonstrate that the subcortical network solution captures the well-established thalamic nuclei configuration. Two established thalamic structures are the lateral geniculate nucleus (LGN), which receives initial visual inputs from the retina via the optic nerve and projects in an organized anatomical fashion to V1 in the mammalian neocortex; and the medial geniculate nucleus (MGN), which relays auditory information from the inferior colliculus to the auditory cortex. Therefore, we tested whether our thalamic solution included the LGN and MGN (**Fig. 6a**). As expected, we observed that the LGN was assigned to the primary visual network (VIS1) and that the MGN was assigned to the auditory network (AUD). Notably, the MGN was not assigned to the auditory network in the woGSR version of the partition (**Fig. 6b**). Structural and functional connectivity of the LGN (**Fig. 6d, f-j**) and MGN (**Fig. 6e, k-o**) parcels defined by the partition reveal connectivity with expected visual (e.g. superior colliculi, primary visual cortex) and auditory (e.g. inferior colliculi, primary auditory cortex) processing regions respectively.

**Figure 6.**
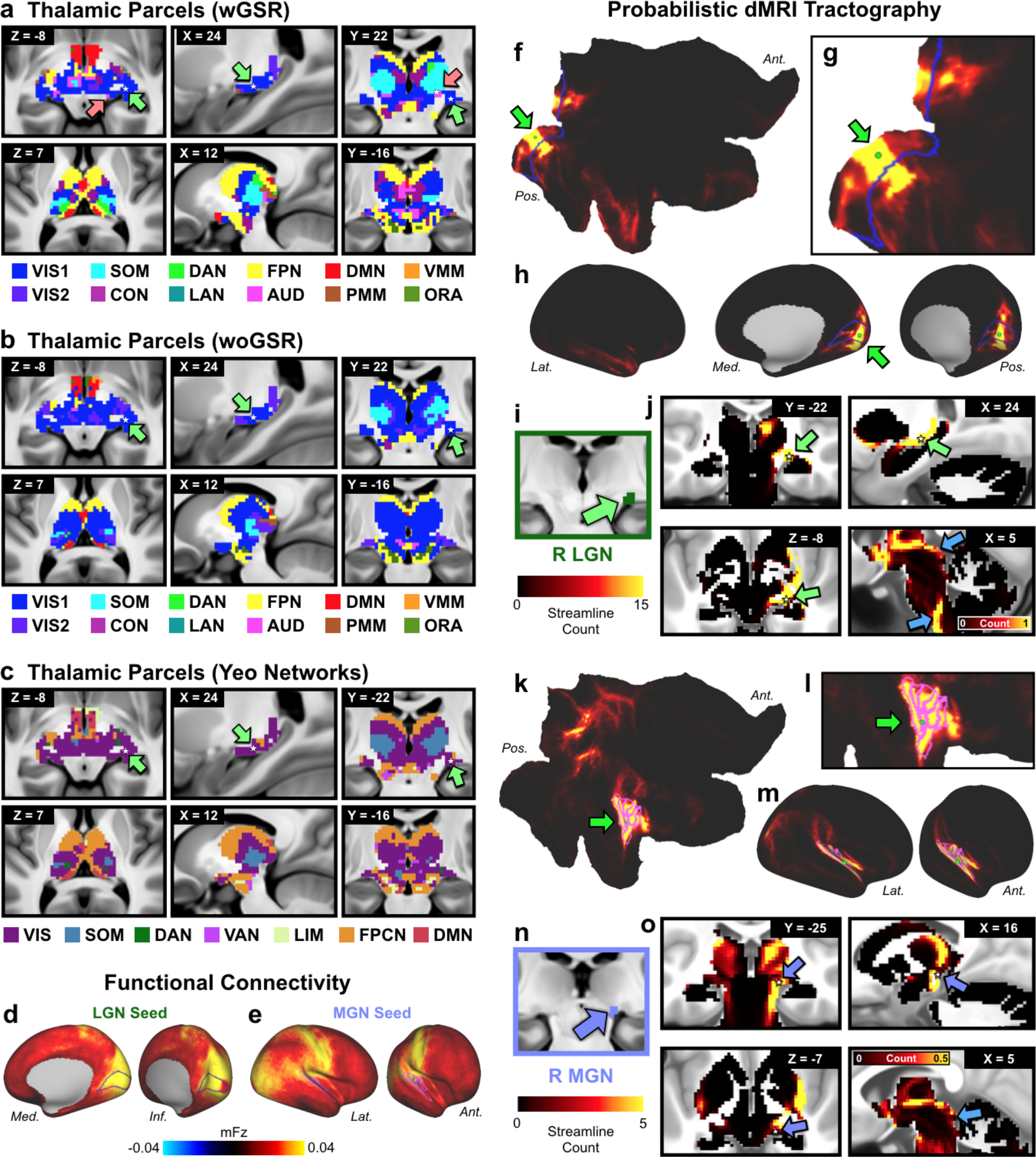
Thalamic network assignment. **A)** Network assignment of the thalamus and ventral diencephalon from the network partition described in the manuscript. Top row highlights the horizontal, sagittal, and coronal views of the lateral geniculate nucleus (LGN), indicated by green arrows, and the medial geniculate nucleus (MGN), indicated by pink arrows. White stars mark the voxel seeded for functional connectivity in D and E. Bottom row shows cross-sectional view of the parcellation at different slices. **B)** Network assignment of the thalamus and ventral diencephalon from the parcellation performed without GSR (woGSR). Without GSR, the auditory network assignment of the MGN was not distinguishable in the parcellation. **C)** Network assignment of thalamus and ventral diencephalon using cortical network parcellation from Yeo et al. (2011). Note the lack of an auditory network in the Yeo et al. (2011) partition limits the ability to map thalamus relative to the new partition reported here. **D)** Cortical functional connectivity of the bilateral LGN parcels. VIS1 parcels are outlined in blue. Right hemisphere is shown; similar results were seen in the left hemisphere. **E)** Cortical functional connectivity of the bilateral MGN parcels. AUD parcels are outlined in blue. Right hemisphere is shown; similar results were seen in the left hemisphere. **F)** Probabilistic tractography (i.e. ‘structural’ connectivity) of the right primary visual cortex (V1) shown in flat cortical map. Seed grayordinate is highlighted with green dot and arrow. Cortical VIS1 network parcels are outlined in blue. Tractography results were computed from diffusion MRI data obtained from the same subjects and averaged over the entire group. **G)** Magnified view of V1 seed (green dot) in flat cortical map. **H)** Inflated cortical view of V1-seeded probabilistic tractography results. **I)** Right LGN identified using the Jülich atlas (Bürgel et al., 2006; Eickhoff et al., 2005), similar coordinates also reported in (Linzenbold et al., 2011; Marx et al., 2004; A. T. Smith et al., 2009). **J)** Tractography of V1 seed to subcortex, including the right LGN (green arrows). White stars mark the right LGN voxel from which functional connectivity was seeded in **D**. Connectivity was strongest between V1, right LGN, and other visual processing regions, including the superior colliculus and brainstem nuclei (blue arrows). Results were similar for the left LGN. **K)** Probabilistic tractography of the right primary auditory cortex, displayed in flat cortical map. Seed grayordinate is highlighted with green dot and arrow. Cortical AUD network parcels are outlined in fuchsia. **L)** Magnified view of primary auditory seed (green dot) in flat cortical map. **M)** Inflated cortical view of auditory-seeded probabilistic tractography results. **N)** Right MGN identified using the Jülich atlas. **O)** Tractography of primary auditory seed to subcortex, including right MGN (purple arrows). White stars mark the right MGN voxel from which functional connectivity was seeded in **E**. Connectivity was strongest between right auditory cortex, right MGN, other thalamic nuclei, and auditory processing regions such as the inferior colliculi (blue arrow). Results were similar for the left MGN. Abbreviations: Lat., lateral; Med., medial; Ant., anterior; Pos., posterior.

### Identification of Novel Functional Networks: Posterior Multimodal, Ventral Multimodal, and Orbito-Affective Networks

Three networks emerged from the reported network detection approach that, to our knowledge, do not correspond to previously-described large-scale networks in the human brain (**Fig. 7**). These networks include PMM (posterior multimodal), VMM (ventral multimodal), and ORA (orbito-affective) networks. We found converging evidence to support the robustness of all three networks. First, all three networks were present for both groups of subjects in the cortical split-half analysis (**Fig. 2c**). Second, all three networks had subcortical representations which were statistically significant at *p*<0.05 (with the exception of VMM) in a split-half replication of those assignments (**Fig. 4d-e**, **Fig. 7**, **Methods**). Third, the PMM and VMM networks were within one standard deviation of the cross-network mean confidence scores, suggesting equivalent confidence in these networks as better-established networks. While the ORA network exhibited the lowest confidence score, it was still well above chance, consistent with ORA regions having higher RSFC among themselves than with regions of other networks. Fourth, the inter-subject variability across the PMM network RSFC patterns was near the mean value across all networks (PMM inter-subject variability=0.41, cross-network mean=0.42), suggesting that PPM inter-subject variability was not appreciably different. In contrast, the VMM and ORA networks had somewhat higher inter-subject variability than the cross-network mean. While it is not possible to assess statistical significance of this result (due to this statistic being calculated across all subjects simultaneously, precluding the ability to use, e.g., t-tests), the high symmetry and replicability of these networks suggest these networks are well-defined. While we cannot altogether exclude the possibility that some of the subcortical network assignments were partially influenced by data smoothing (particularly in the case of VMM), together these results suggest that the three novel network identified here are robust and are therefore likely to be of broad functional relevance. It will nonetheless be important for future studies to further validate the existence of these network and better determine their functional roles.

**Figure 7.**
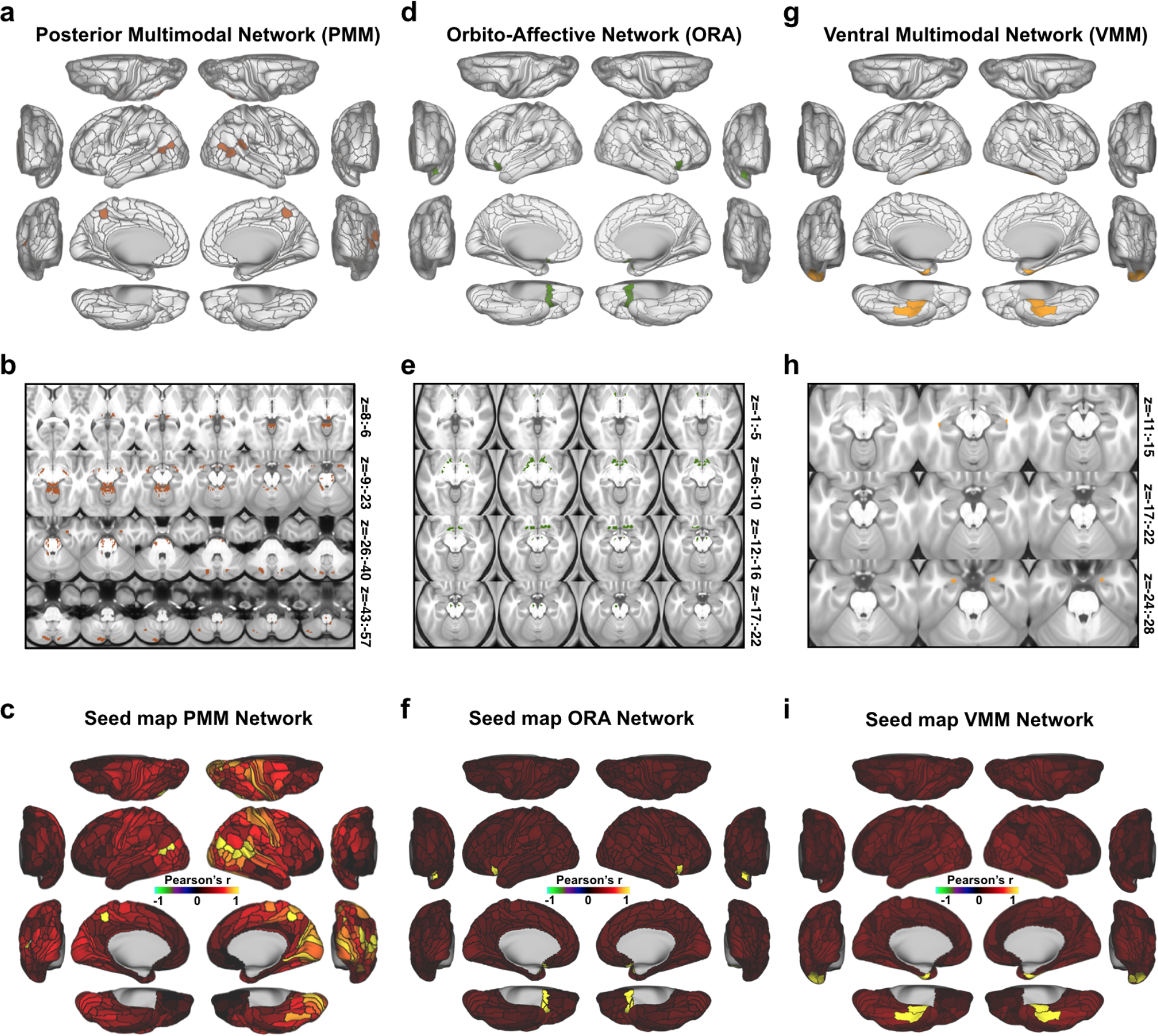
New posterior multimodal, orbito-affective, and ventral multimodal networks. **A)** Cortical parcels that are part of the posterior multimodal (PMM) network as detected by the Louvain clustering algorithm. **B)** Subcortical areas that were identified as PMM based on correlation with cortical regions. **C)** Cortical seed map of the PMM network showing connectivity to all other parcels (within-network connectivity is 1 in all PMM parcels). **D)** Cortical parcels that make up the orbito-affective (ORA) network as detected by the Louvain clustering algorithm. **E)** Subcortical areas associated with the ORA network. **F)** Cortical seed map of the ORA network showing connectivity of this network to all other cortical parcels.**G)** Cortical parcels that are part of the ventral multimodal (VMM) network as detected by the Louvain clustering algorithm. **H)** Subcortical areas associated with the VMM network. **I)** Cortical seed map showing connectivity of the VMM network to all other parcels.

### Characterizing the Laterality and Function of the Language Network

As mentioned above, the network identified as a language network (LAN, including well-known language-related areas such as Broca’s and Wernicke’s areas) showed high asymmetry for its regions’ cortex-wide RSFC patterns. To further test the hypothesis that this network carries out language-related functionality, we first analyzed the LANGUAGE task fMRI data provided by the HCP to map the amount of overlap of the derived whole-brain LAN network with language-activated grayordinates (see **Fig. 8a-b** for cortical and subcortical maps). This overlap was significantly higher than expected by chance (**Fig. 8c**), suggesting that these areas are indeed largely overlapping with language processing areas (>85% observed overlap). Second, we quantified the network’s asymmetry (see **Methods**) by calculating asymmetry for each cortical parcel (**Fig. 8d**) and subcortical voxel (**Fig. 8e-f**). Compared to other networks, the LAN network was appreciably more asymmetric in cortex (LAN vs VMM: t(336)=3.38, *p*=0.0008, LAN vs. mean of all other networks: *p*<0.00001, also see **Fig. 8e**). Further, there were more LAN parcels identified in the left hemisphere (14 parcels) than the right hemisphere (9 parcels) of cortex. Also in subcortex, LAN emerged as one of the most asymmetric networks, as can be seen when comparing the proportion of non-overlapping subcortical voxels in left and right hemispheres. Similar left lateralization as in cortex was observed in subcortex when quantifying the proportion of total voxels in left and right hemisphere (left and right reversed for cerebellum, as expected). This asymmetry far exceeded chance levels (chance proportion of voxels in left subcortex/right cerebellum for all networks=0.50; proportion of voxels in left subcortex/right cerebellum for LAN=0.71;*χ*^2^=8.878, *p*=0.0029). In turn, we focused on a single asymmetric left-lateralized LAN region, area PSL. RSFC seed maps of left and right PSL (**Fig. 8g-h**) were strikingly different, with left PSL showing high LAN connectivity and low CON connectivity, but right PSL showing the opposite pattern. The LAN regions overlapping with language task activations, observed strong left-lateralized lateralization, and qualitatively-distinct connectivity patterns in asymmetric regions together strongly support the hypothesis that this network implements language functionality.

**Figure 8.**
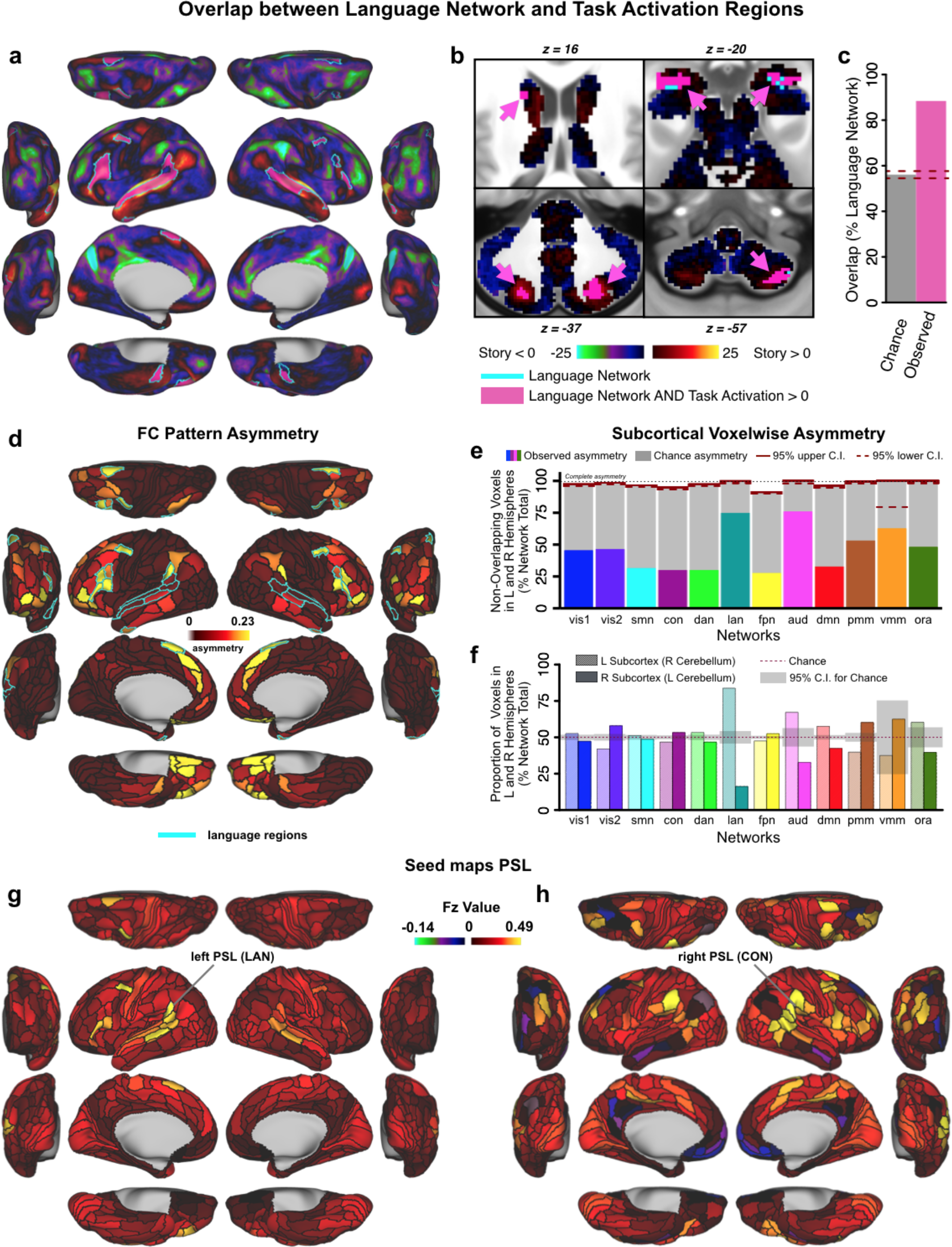
Language network evaluation. **A & B**)Overlap between the language network (LAN, teal outline) from our resting-state based network partition and activations from an independent language processing task (collected in the same sample of 337 subjects) in cortical and subcortical regions. Pink areas indicate overlap between LAN and task activation. Underlay shows task activation t-statistics from the ‘Story versus Baseline’ contrast of the LANGUAGE task, replicating the analysis conducted by Glasser and colleagues (2016). Note that t-scores are shown here because the high statistics resulted in infinity values when converting to Z-scores. **C)**Percentage overlap between LAN and task activation in the language processing task expected by chance (gray bar) and actual observed overlap in panels A&B (pink bar). Dashed lines indicate 95% confidence intervals. **D)** Cortical map displaying the asymmetry of parcels. The teal outline indicates the language network, which is highly asymmetric compared to the other networks, with left hemisphere dominance. **E & F)** Network asymmetry in the subcortex. Colored bars in Panel E show the proportion of subcortical voxels in each network that do not overlap when comparing left and right hemispheres. Complete asymmetry (no overlap) is indicated by dotted line at 100% for reference. gray bars indicate chance asymmetry calculated given the size of each network. Solid and dashed red lines indicate 95% upper and lower confidence intervals for chance respectively. Panel F displays the proportion of total voxels in left and right hemispheres for each network. Chance level for this measure is 50% for all networks; confidence intervals are calculated given the total number of voxels in each network. Because functional representation of left and right is reversed in the cerebellum relative to the rest of the brain (due to the midline crossing of projecting fibers (van Baarsen et al., 2016)), left and right cerebellar hemispheres were exchanged in calculating this measure. Like the cortical networks, panel E&F show that subcortical networks are symmetric in general, with a left lateralized LAN. In subcortex, VMM is also significantly asymmetric. **G & H)** Functional connectivity seed maps for left and right perisylvian language areas (PSL) based on resting-state data in 337 subjects. Both the left and right language seed area show strongest connections to ipsilateral regions.

### Improved Reproducibility and Statistics of Language-Related Activation Using the Cortical-Subcortical Network Partition

We next sought to demonstrate the practical utility of the network partition and its beneficial impact on actual data analysis. The partition could be applied in a variety of ways, such as interpreting task-evoked activations or functional connections in terms of a canonical set of functional networks. For this demonstration we focused on the identification of a putative “language” network. If this mapping is veridical in relation to the language system, then we hypothesized two effects to emerge: i) There should be high overlap between the language network and the task-evoked signal produced by the ‘Story versus Baseline’ LANGUAGE task (demonstrated in **Fig. 8**); ii) There should be an appreciable statistical improvement in the ‘Story versus Baseline’ LANGUAGE task contrast when going from a ‘dense’ grayordinate-level effect to a parcellated effect (as shown for several language-related local areas by Glasser et al. (2016)) in language network regions. Additionally, if the identified language network indeed maps onto independently-defined language-related task-evoked fMRI signal, then there should be even greater statistical improvement if computing the GLM-derived task-evoked signal across the entire language network. Showing such a statistical improvement would demonstrate a powerful and empirically useful application of the network partition for detecting neurocognitive effects in a more robust way.

To address the second hypothesis, we calculated statistics for the ‘Story versus Baseline’ LANGUAGE task contrast after separately fitting the task GLM to: dense grayordinate-level time series data (**Fig. 9a-b**, identical to underlay in **Fig. 8a**), time series data averaged within a given parcel (**Fig. 9c-d**), and time series data averaged within a given whole-brain network (**Fig. 9e-f**). As hypothesized, the t-statistic across the whole-brain LAN network was markedly higher when data were first averaged at the network level before fitting the task GLM, compared to fitting the task GLM on the ‘dense’ grayordinate level or parcel-level time series and then averaging across LAN regions (network t=24.71; parcel mean t=12.38, SD=10.91; dense mean t=8.50, SD=6.98; **Fig. 9g**). This effect was robustly present within the cortex (network t=24.71; parcel mean t=10.93, SE=11.66; dense mean t=8.74, SD=7.00; **Fig. 9h**) and the subcortex (network t=24.71; parcel mean t=1.45, SD=3.86; dense mean t=3.37, SD=3.72; **Fig. 9i**). Overall, t-statistics for all three LANGUAGE task contrasts were markedly improved by fitting the task GLM to parcel-level time series, rather than fitting to dense time series and averaging across parcels afterwards (**Fig. 9j**, note sigmoidal deviation from diagonal). Importantly, t-statistics were further improved when the task GLM was fit on network-averaged time series, compared to parcellating by network after fitting on dense (**Fig. 9i**) or parcel time series (**Fig. 9l**). This result strongly supports that the signal-to-noise ratio was substantially improved by first averaging BOLD time series data within the identified LAN network. Of note, this result also reinforces the inference that the LAN asymmetry reflects true lateralization. Together, these task-evoked effects add confidence to the identified language network definition and demonstrate the practical utility of the network partition, which is released publicly as part of this study.

**Figure 9:**
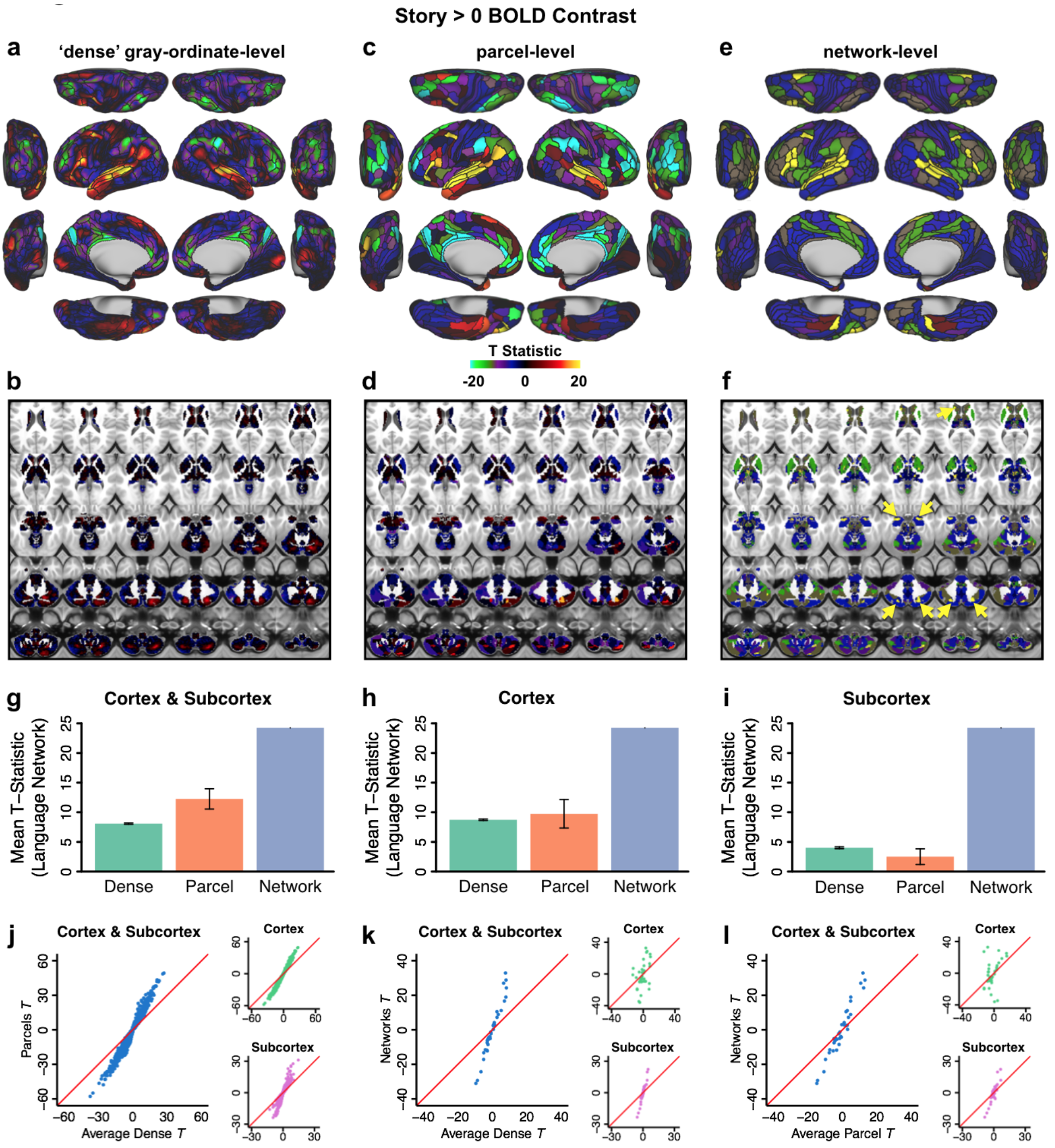
Demonstration of improved reproducibility and statistics with new partition. Panels A-F show task activations for a language processing (LANGUAGE task ‘Story versus Baseline’ contrast) task at three different levels. **A)**Cortical activation map of dense-level analysis. **B)**Subcortical activation map of dense-level analysis. **C)**Cortical activation map of parcel-level analysis. Task fMRI data were first parcellated at the parcel level before model fitting. **D)**Subcortical activation map of parcel-level analysis. **E)**Cortical activation map of network-level analysis. Task fMRI data were first parcellated at the network level before model fitting. **F)** Subcortical activation map of network-level analysis. Yellow arrows highlight subcortical regions with a high task-activated t-score, which overlap with parcels in the LAN network. **G)** t-statistics (LANGUAGE task ‘Story versus Baseline’ contrast) shown in panels A-F significantly improve for the parcel-level vs. dense-level analysis, and for the network-level vs. parcel-level analysis in a combined cortex and subcortex analysis. Error bars are inter-parcel standard deviations. **H)** t-statistics (LANGUAGE task ‘Story versus Baseline’ contrast) in cortex alone again show significantly better results for the network-level analysis compared to the dense-and parcel-level analyses. **I)** t-statistics (LANGUAGE task ‘Story versus Baseline’ contrast) in subcortex showed substantially better results for the network-level analysis compared to the dense-and parcel-level analyses. Note that – in contrast to the results for cortex – parcel-level analysis in subcortex does not give an advantage over dense-level analysis. **J)** An improvement in t-statistics was found when task designs were fit on parcellated time series instead of on dense time series and subsequently averaging for parcels. Blue dots represent 718 parcels × 3 LANGUAGE task contrasts (‘Story versus Baseline’; ‘Math versus Baseline’; ‘Story versus Math’). Insets show the 360 cortical parcels × 3 contrasts (top, green dots) and 358 subcortical parcels × 3 contrasts (bottom, purple dots) separately. **K)** Improvement in t-statistics was also found when fitting task designs on network time series compared to fitting on dense time series and then averaging for networks. Blue dots represent 12 networks × 3 LANGUAGE task contrasts. Insets show the 12 cortical networks × 3 contrasts (top, green dots) and 12 subcortical parcels × 3 contrasts (bottom, purple dots) separately. **L)** A further improvement in t-statistics was found when fitting on networks versus fitting on parcels and then averaging for networks. Blue dots represent 12 networks × 3 LANGUAGE task contrasts. Insets show the 12 cortical networks × 3 contrasts (top, green dots) and 12 subcortical parcels × 3 contrasts (bottom, purple dots) separately.

## DISCUSSION

The human brain is a unified dynamical computational system that, ultimately, can only be understood as a whole. Simultaneously, understanding any dynamical system requires identifying its functional components and their interactions. We therefore sought to build on previously-developed network partitions to create a whole-brain network partition, identifying large-scale network communities of brain regions across both cortex and, for the first time, all subcortical areas. We created this whole-brain partition as a resource to aid neuroscience research generally, and we are therefore releasing the partition (along with the data and code that produced it) to the neuroscience community (available at https://github.com/ColeLab/ColeAnticevicNetPartition once through peer review).

As with all neuroscientific methods there are limitations to the approach used here (detailed below), but also several advantages. First, we used a large dataset relative to most neuroscientific studies to date (337 subjects), increasing the effective SNR and the likelihood that the results will generalize to new groups of individuals. Second, we used multiple quality control metrics to ensure stability and reliability of the network partition, which were found to be fairly high by all applied standards. Third, we used a principled approach to decide on the network partition algorithm and associated parameters, involving both stability optimization and calibration of parameters based on well-established neurobiologically-grounded constraints (e.g. the existence of primary sensory-motor networks). Fourth, we extended the cortical network partition to subcortical structures, resulting in a comprehensive map of brain-wide functional networks. Finally, we used task fMRI data to demonstrate a practical advantage of using this network partition: an increased ability to detect network-level functional activations.

Notably, previously published large-scale network partitions have already made a substantial positive impact on neuroscientific investigations across health and disease. We expect the network partition developed here to also be useful across a variety of neuroscientific investigations. For instance, the network partition could be used to interpret possible functions of a region-level activation using fMRI, EEG, or local field potentials. Alternatively, the network partition could be used in studies of network dynamics, placing interactions in a larger functional context to aid in summarizing and interpreting results. Another use of the network partition could be as a data reduction approach, increasing data processing efficiency while maintaining functionally-meaningful large-scale network units. Finally, this partition makes it possible to test hypotheses about brain-wide functional network organization, spanning cortex, striatum, thalamus, amygdala, hippocampus, brainstem, the cerebellum, and other structures. It is also notable that unlike the brain region and brain network levels, lower levels of organization such as the neuron or local circuit are not expected to generalize across individuals. This is due to the very low likelihood of functionally-equivalent individual neurons aligning anatomically between individuals. Thus, like identifying brain regions, characterizing large-scale brain networks provides units of brain organization that can provide a testbed for the following question, “what does this brain structure do functionally across individuals?” – a key question for generalized understanding of human brain function.

### Extending Prior Network Partitions to Converge on a Global Characterization of Human Brain Network Organization

The network partition identified here is, as expected, similar in many ways to previously-identified network partitions. However, there are several differences that provide novel discoveries regarding the large-scale network architecture of the human brain. Similar to both the Power et al. (2011) and Yeo et al. (2011) cortical network partitions, a variety of well-known sensory-motor and previously-discovered cognitive large-scale functional networks were identified. Common to both of these network partitions, we identified FPN, CON, DMN, DAN, visual, and somatomotor networks. Unlike Yeo et al. but similar to Power et al., we identified a separate auditory network consistent with the primary auditory cortical system. Notably, this auditory network was merged with the somatomotor network at various parameter settings of our network detection algorithm, consistent with the Yeo et al. result. This illustrates the difficulty of identifying the correct “data-driven” metrics when using a clustering algorithm – auditory and somatomotor regions are known to perform highly distinct functions yet their RSFC patterns were difficult to separate without explicitly forcing this neurobiologically-established separation. We identified three networks that, to our knowledge, have not been previously identified:

PMM (posterior multimodal), VMM (ventral multimodal), and ORA (orbito-affective). Unlike the language network, we did not predict the existence of these networks based on the prior literature. Importantly, lack of pre-existing evidence of the VMM and ORA networks was likely driven by signal dropout in the proximity of these networks, due to magnetic field inhomogeneities from nearby sinuses (Deichmann et al., 2003). The multiband fMRI data used here (Uğurbil et al., 2013) appears to have reduced the signal dropout near sinuses. This is likely due to smaller voxels (2 mm cubic voxels used here rather than the standard-to-date 3+ mm voxels), which can reduce MRI signal dropout (Merboldt et al., 2000; Smith et al., 2013). The increased precision from using a cortical surface analysis (Anticevic et al., 2008), averaging across vertices within each parcel, and averaging with a large sample size likely all contributed to an increased ability to map RSFC in these dropout areas. While we found some evidence for lower SNR in these regions relative to other cortical regions, we also identified strong reliability of the networks using split-half validation (**Fig. 2c** & **Fig. 3d**) and found a symmetric, replicable and robust subcortical contribution to these networks, further validating these networks. It will be important for future studies to corroborate the existence of these networks, identify their functional roles, and enumerate the factors (such as voxel size) that affect the ability to detect them.

It is unclear at this point what functions these networks might perform, given that they represent a novel discovery. While we appreciate this partially reflects reverse inference, we used observations of the constituent parts of the networks to infer possible functionality and provide a label. This is most evident in the case of the ORA network, which overlaps with portions of cortex associated with “reward processing” functionality in posterior orbitofrontal cortex (Kahnt et al., 2011; Schultz, 2006). Corroborating this interpretation, ORA connected strongly with known reward-related areas in subcortex. These included the ventral striatum (Delgado et al., 2000; Schultz et al., 1992), midbrain nuclei consistent with the substantia nigra / ventral tegmental area (which contain dopamine neurons) (Fiorillo et al., 2003), and the globus pallidus (Justin Rossi et al., 2017). Further, this portion of cortex was modulated differentially by rewarding stimuli (Camara et al., 2010). This is consistent with a strong role for reward-related dopamine projections to ORA, suggesting strong reward processing functionality for this network.

The VMM network consists of four cortical regions on the ventral surface of the temporal lobe. The VMM extends into subcortex only minimally, with a cluster in the right ventral striatum and small bilateral clusters in the hippocampus. One possible function of this network is to represent higher-order semantic categories, consistent with studies of anterior (Rogers et al., 2006) and inferior temporal lobe (De Baene and Vogels, 2010). The novelty of this network reduces our ability to identify its functionality, however, such that it will be important for future studies to better determine what the functional specializations of this network might be.

The PMM network consists of bilateral dorsomedial parietal lobe, bilateral temporal-parietal-occipital junction, and right dorsocaudal temporal lobe. The PMM also extends into a variety of subcortical locations. These locations include: bilateral amygdala, portions of the brainstem, the putamen, multiple portions of cerebellum, a small portion of the caudate, a small portion of thalamus, and a portion of the diencephalon consistent with the basal forebrain. Most of these subcortical locations were assigned symmetrically across hemispheres and showed strong split-half replication. This demonstrates that while these assignments were widespread they were nonetheless robust, suggesting the existence of previously-unknown widespread PMM circuits. One possible function of this network could be spatial navigation, given the importance of dorsomedial parietal lobe for spatial navigation (Marchette et al., 2014). Additionally, PMM might be important for identifying and representing event structure in narratives, given that PMM regions were recently shown to represent long narrative structures during movie watching (Baldassano et al., 2017). It will be important for future studies to carefully map the PMM as identified here to particular functions such as spatial navigation and representing situational/narrative structures.

Our ability to map the ventral surface of cortex presented a unique opportunity, since most previous network partitions omitted these territories due to MRI signal dropout. We not only identified two novel networks in these dropout zones, but were also able to test for expansion of previously-identified networks into an extensive portion of cortex for the first time. We found that orbitofrontal cortex (OFC) was split into thirds, with nearly equal assignment of OFC parcels to FPN, DMN, and ORA. It is notable that so much of OFC was assigned to FPN given that the FPN is classically described as primarily lateral prefrontal cortex and parietal cortex. This result suggests that the task-rule-oriented representations in lateral prefrontal cortex (Cole et al., 2011; Stokes et al., 2013) likely interact extensively with action-outcome and stimulus-reward associations in OFC (Kahnt et al., 2011). Indeed, some nonhuman primate studies have suggested such interactions occur during task performance (Wallis and Miller, 2003). The present study suggests these interactions occur as a part of a global system likely specialized for cognitive control and associated goal pursuit (Cole et al., 2014b, 2013; Duncan,2010). It will be important for future studies to more fully characterize the relationships between classic portions of FPN and these portions of OFC previously unassigned due to MRI signal dropout.

### Mapping a left-lateralized brain-wide language network in the human brain

Unlike many previous network partitions, we identified a whole-brain network highly consistent with language functionality. This was based on the proximity of its regions to the well-established Broca’s and Wernicke’s areas, its left lateralization being consistent with known left lateralization of language functionality (Gazzaniga, 2005; Gazzaniga et al., 1962), as well as its activation during a language task. Additionally, several of the regions included in this network were thoroughly investigated by Glasser et al. (2016), establishing these regions as distinct functional entities with clear language functionality. Notably, the Power et al. (2011) partition (updated and visualized more fully by (Cole et al., 2013)) included a network consistent with this language network, but labeled the “ventral attention network”. The present results suggest this network was previously mislabeled, since its connectivity pattern, anatomical location, and task activations are most consistent with language functionality.

One key feature of the language network identified here is its left lateralization. We found that the cortical language network was the most lateralized network in terms of RSFC pattern asymmetry (**Fig. 3e**, **8g, & 8h**), that the subcortical voxels assigned to the language network were more left-lateralized than expected by chance (**Fig. 8e & 8f**), and that the language network overlapped more with language task activations than chance (**Fig. 8a, 8b & 8c**). Lateralization of language functionality is one of the most well-established findings in the human brain (Mesulam, 1998), making it somewhat surprising that this has not been emphasized in previous RSFC literature. A recent study (McAvoy et al., 2015) found that left-lateralized language functionality only emerged in their RSFC analysis when global signal regression was not included as a preprocessing step. Inconsistent with this, however, the Power et al. (2011) network similar to our identified language network was left lateralized (with dorsal and medial frontal network assignments being more extensive in the left hemisphere), despite use of global signal regression. Thus, while not performing global signal regression may have assisted our identification of the language network, it was unlikely that avoiding global signal regression was necessary to identify this network.

Beyond simply counting more language-assigned parcels in the left hemisphere (14 left, 9 right), our use of pattern asymmetry was important for precisely quantifying lateralization. RSFC pattern asymmetry revealed that several language network regions had highly distinct global patterns of RSFC with their right-hemisphere homologues. This striking qualitative difference across homologous parcels is illustrated in detail in **Fig. 8g & 8h**. This parcel, which is consistent with Wernicke’s area on the left, was assigned with high confidence to the language network on the left but with high confidence to CON on the right. Consistent with this assignment difference, many regions with low RSFC for the left hemisphere parcel are high for the right hemisphere parcel, and vice versa. Together these results demonstrate the strength of left lateralization of the language network, both in terms of the number of left-lateralized parcels, asymmetry of global RSFC patterns, as well as its subcortical contributions.

### Mapping the complex relationships between subcortical structures and cortical networks

We found that all 12 cortical functional networks, including higher-order associative networks (such as the FPN and CON), were reliably represented across the entire subcortex and the cerebellum. This is consistent with known functional loops between all portions of cortex and thalamus (Behrens et al., 2003), which in turn loop through basal ganglia (Middleton and Strick, 1994) and cerebellum (Kelly and Strick, 2003). Also consistent with the observed widespread connectivity between cortex and subcortical nuclei, various subcortical nuclei involving a variety of neurotransmitters (e.g., substantia nigra, basal forebrain, raphe nucleus) are known to project broadly throughout cortex (Herlenius and Lagercrantz, 2004). Finally, regions such as amygdala (Barbas, 2000; Jolkkonen and Pitkänen, 1998) and hippocampus (Eichenbaum et al., 2007) are thought to project to and from multiple cortical networks. Importantly, most of what is known about these subcortical structures comes from non-human animal studies or localized functional neuroimaging studies in humans, with relatively few focused RSFC studies (Buckner et al., 2011; Choi et al., 2012). The reported results represent the first comprehensive attempt to assign each subcortical voxel to a given cortical network. In turn, we establish the replicability, stability, symmetry and task-evoked relevance of such a subcortical functional network solution. Nevertheless, there were some unexpected findings that will be important to follow up on in future research. First, we found that the language network exhibits notable connectivity with the amygdala. Second, we identified a large and robust subcortical contribution to the primary visual network, perhaps reflecting a distributed ‘attentional system’, involved in overt attention and wakefulness. Of note, we did explicitly enforce a separate of the V1 and secondary visual cortical networks. Recent work suggests that there may be some residual artifact associated primarily with visual and somatomotor systems (respiration, sleep, movement) (Bijsterbosch et al., 2017; Glasser et al., 2017). It may be possible that assignment of some subcortical structures to VIS1 is inflated due to this artifact (perhaps due to eyes open vs. closed correlating with sleep+respiration changes). This current limitation that can be improved in future iterations of the partition by leveraging recently proposed advances in temporal de-noising that circumvents global signal removal (Glasser et al., 2017).

Importantly, prior subcortical network assignment attempts did not incorporate the thalamus and the brain stem in their reported solutions. As noted, thalamic sub-nuclei are well-known to form functional circuits with cortical networks (Barbas, 2000; Zhang et al., 2008) and have been shown to exhibit robust patterns of diffusion MRI-derived probabilistic tractography with cortical territories (Behrens et al., 2003). Therefore, it was vital to demonstrate that the subcortical network solution captures the well-established thalamic nuclei configuration. A ubiquitously established thalamic structure is the lateral geniculate nucleus (LGN), which receives initial visual inputs from the retina via the optic nerve and projects in an organized anatomical fashion to V1 in the mammalian neocortex. Therefore, we established that our thalamic solution included LGN. We observed a well-preserved correspondence of the thalamic network assignment whereby the LGN was encompassed by the primary visual network (VIS1) (see **Fig. 6**).

### Limitations and opportunities for further improvement of the network partition

There are several limitations to the approach used here that represent important opportunities for future improvements to understanding the large-scale functional organization of the human brain. For instance, any network partition necessarily oversimplifies brain organization by removing/downplaying inter-network interactions. Nonetheless, it is useful to know the overall network organization while acknowledging the smaller/rarer interactions between networks. Additionally, this is not a fully exhaustive search over all possible network organizations. Our partition was likely not fully optimal due to the need to use heuristics to identify network organization (for computational tractability) (Blondel et al., 2008; Girvan and Newman, 2002). This leaves open the possibility of more accurate network organizations in the future. Nonetheless, we assessed multiple algorithms and ran a large-scale parameter search, achieving a highly optimal and reliable network partition as quantified by a variety of quality assessment metrics.

Despite covering the whole brain (unlike most previous network partitions), we nonetheless maintained a cortical-centric approach. Specifically, we began by creating a cortical network partition, which was then extended into subcortical voxels by quantifying the relationship between subcortex with the cortical networks. This may introduce a cortico-centric bias as the subcortical solution is explicitly driven by defining the cortex partition first. Nonetheless, we used this approach to aid in bridging the currently cortico-centric view of human brain function to subcortical structures. We also used this approach given the historical utility of understanding subcortical functions based on connectivity with specific cortical structures. For instance, mapping cerebellar connectivity with cortex in macaque monkeys has aided in understanding functional specialization in cerebellum (Kelly and Strick, 2003). Furthermore, this approach has proved highly productive and impactful in prior attempts at mapping striatum and cerebellum onto cortical networks (Buckner et al., 2011; Choi et al., 2012). We nonetheless expect that future research will benefit from a more even-handed partitioning of cortical-subcortical gray matter. This would involve creating functional-defined three-dimensional brain parcels in subcortical structures, just as was done as an initial step in cortex (Glasser et al., 2016). These parcels would then be included in a community detection algorithm along with the cortical parcels. This may reveal distinct subcortical parcels from what we identified here, in addition to potentially distinct networks. Notably, it is possible that two functional parcels that are neighbors in anatomical space could be merged in our current approach if they were both assigned to the same network. Nonetheless, we expect that our approach has advanced understanding of subcortical structures, putting them in the functional context defined by large-scale cortical networks.

Additional improvements on the present network partition could stem from even more precisely defining the cortical parcels. Presently used parcels were identified based on convergence across multiple neuroimaging modalities (e.g., fMRI and structural MRI), likely limiting biases from any one modality. Nevertheless, certain decisions were made when deriving this parcellation that may be reconsidered in future. For instance, the Glasser parcels force the face and non-face representations in the primary motor homunculus to be merged (since primary sensory and primary motor regions were defined in part based on cytoarchitecture), even though it is clear that these portions of the motor homunculus have distinct RSFC patterns (Power et al., 2011; Yeo et al., 2011). Despite such potential limitations the use of multiple modalities when defining parcels by Glasser and colleagues likely reduced biases present in any one modality (e.g., RSFC).

There is evidence that global signal removal (GSR) is important for reducing respiratory and motion artifacts that plague RSFC (Power et al., 2017b, 2014). GSR was not used for the primary analyses in the cortex of the current study because Glasser et al.(2016) reported that GSR appreciably shifts RSFC gradients (relative to other modalities only minimally affected by respiratory/motion artifacts) used for identifying the cortical parcels, which could invalidate use of these regions in the present study. However, we used GSR in the subcortical parcellation to test the hypothesis that GSR could reduce the extensive assignment of low-SNR subcortical voxels to the visual networks (**Supplemental Fig. S3**). As expected, the visual networks were less extensively represented in the version with GSR, but all networks were replicated significantly above chance in both the wGSR and woGSR versions. Because the wGSR version of the subcortical partition revealed known neurobiological structures such as the MGN as well as higher convergence of SMN with motor task activation, we present this version of the subcortical parcellation in our primary analyses, although many of the analyses using the woGSR version revealed comparable results (**Supplemental Fig. S3-S4**). Importantly, GSR may serve to reduce artifact-related noise in particular in the subcortex of these HCP data. Other studies in the literature have demonstrated that GSR helps to reduce noise even in data that has undergone ICA-FIX (Power et al., 2017a, 2017b, 2014), suggesting a need for further improvements in methods for removing global noise. Simultaneous with the present study an approach involving temporal independent components analysis (ICA) has been developed to remove global noise while leaving global signal of neural origin (Glasser et al., 2017). This results in RSFC with global noise distortions removed without GSR-driven distortions such as RSFC gradient shifts. Future work should generate a revised version of the partition after the parcellation has been re-computed using this new temporal ICA de-noising method.

We identified several previously-unidentified networks, finding at they replicated across independent sets of subjects. A major limitation of these discoveries, however, is the possibility that noise properties particular to the MRI sequences and scanner biased the results, driving the observed replication. It would be helpful to use alternative MRI sequences and scanners (or even highly distinct methods such as magnetoencephalography) to rule out this possibility in order to better validate these new networks. Additionally, these networks would be better validated if they were found to match coactivation patterns using task fMRI, as was the case with the language network being active during the language task and the somatomotor network during the motor task here.

Another opportunity for future improvement is to better characterize the hierarchical nature of brain network organization. This reflects the fact that network organization is likely hierarchical in the sense that each large-scale brain network could be broken down into smaller and smaller components, eventually reaching the single-region level. Critically, however, we used a principled approach to define our target level of organization by setting parameters to detect well-established primary sensory-motor cortical systems. Thus, we created a whole-brain network partition intentionally defined as being at (or near) the same level of organization as these well-established brain systems. We therefore expect that the calibration of our community detection algorithm likely identified networks in association cortex that are at the same (or a similar) level of organization, and are therefore of similar importance for higher-level cognitive functions as primary sensory-motor systems are for perceptual-motor functions. Notably, this principled calibration may have led some previously-identified networks (such as the “salience” network (Power et al., 2011; Seeley et al., 2007)) to not be identified here, likely because they are at a lower level of organization (e.g., salience network being part of the cingulo-opercular network identified here) than the brain systems used for calibration. It will be important for future network partition efforts to characterize the hierarchical sets of networks at different levels of organization.

### Conclusions

The results presented here describe the current version (v1.0) of a novel whole-brain functional network characterization of the human brain. The primary purpose of this study is to describe the network partition dataset, which is now publicly available. We additionally reported a series of quality assessments and validations of the provided network partition. We found evidence that the partition was of high quality and exhibited robust replicability across independent samples as well as across cortical and subcortical structures. While we propose a number of important future improvements of the provided version 1.0, this constitutes the most accurate estimate of whole-brain functional network organization in humans to date. We additionally demonstrated the existence of novel functional networks, such as the lateralized language network, providing additional understanding of human brain organization. The result was successfully applied to a language fMRI task, demonstrating strikingly improved statistical power to detect task-related activations when using the network partition. Collectively, this study demonstrates the value of this whole-brain network partition for scientific inquiry into human brain organization as well as specific task functionality.

Author Contributions (using CRediT Taxonomy, http://www.cell.com/pb/assets/raw/shared/guidelines/CRediT-taxonomy.pdf)
Conceptualization, K.K., A.A., M.W.C.; Methodology, M.S., J.L.J., K.K., G.R., A.A., M.W.C.; Formal Analysis, M.S., J.L.J., K.K.; Data Curation, J.L.J., K.K.; Visualization, M.S., J.L.J.; Writing – Original Draft, M.S., J.L.J., A.A., M.W.C.; Writing – Review & Editing, M.S., J.L.J., G.R., A.A., M.W.C.; Supervision, A.A., M.W.C.

## Acknowledgements

Data were provided by the Human Connectome Project, WU-Minn Consortium (Principal Investigators: David Van Essen and Kamil Ugurbil; 1U54MH091657) funded by the 16 NIH Institutes and Centers that support the NIH Blueprint for Neuroscience Research; and by the McDonnell Center for Systems Neuroscience at Washington University. This work was supported by the NIH via awards K99/R00-MH096801 (Cole), DP5-OD012109 (Anticevic), R01-MH109520 (Cole), R01-MH108590 (Anticevic), R01-AG055556 (Cole), and R01- MH112189 (Anticevic), as well as the Brain and Behavior Foundation (NARSAD) Independent Investigator grant (Anticevic) and ARRS J7-6829 (Repovs).

## REFERENCES

Anticevic, A., Dierker, D.L., Gillespie, S.K., Repovs, G., Csernansky, J.G., Van Essen, D.C., Barch, D.M., 2008. Comparing Surface-Based and Volume-Based Analyses of Functional Neuroimaging Data in Patients with Schizophrenia. Neuroimage 41, 835–848.

Baldassano, C., Chen, J., Zadbood, A., Pillow, J.W., Hasson, U., Norman, K.A., 2017. Discovering Event Structure in Continuous Narrative Perception and Memory. Neuron 95, 709–721.e5.

Barbas, H., 2000. Connections underlying the synthesis of cognition, memory, and emotion in primate prefrontal cortices. Brain Res. Bull. 52, 319–330.

Barch, D.M., Burgess, G.C., Harms, M.P., Petersen, S.E., Schlaggar, B.L., Corbetta, M., Glasser, M.F., Curtiss, S., Dixit, S., Feldt, C., Nolan, D., Bryant, E., Hartley, T., Footer, O., Bjork, J.M., Poldrack, R., Smith, S., Johansen-Berg, H., Snyder, A.Z., Van Essen, D.C., Consortium, for the WU-Minn HCP, 2013. Function in the human connectome: Task-fMRI and individual differences in behavior. Neuroimage 80, 169–189.

Behrens, T.E.J., Johansen-Berg, H., Woolrich, M.W., Smith, S.M., Wheeler-Kingshott, C.A.M., Boulby, P.A., Barker, G.J., Sillery, E.L., Sheehan, K., Ciccarelli, O., Thompson, A.J., Brady, J.M., Matthews, P.M., 2003. Non-invasive mapping of connections between human thalamus and cortex using diffusion imaging. Nat. Neurosci. 6, 750–757.

Bijsterbosch, J., Harrison, S., Duff, E., Alfaro-Almagro, F., Woolrich, M., Smith, S., 2017. Investigations into within- and between-subject resting-state amplitude variations. Neuroimage 159, 57–69.

Binder, J.R., Gross, W.L., Allendorfer, J.B., Bonilha, L., Chapin, J., Edwards, J.C., Grabowski, T.J., Langfitt, J.T., Loring, D.W., Lowe, M.J., Koenig, K., Morgan, P.S., Ojemann, J.G., Rorden, C., Szaflarski, J.P., Tivarus, M.E., Weaver, K.E., 2011. Mapping anterior temporal lobe language areas with fMRI: a multicenter normative study. Neuroimage 54, 1465–1475.

Biswal, B., Yetkin, F.Z., Haughton, V.M., Hyde, J.S., 1995. Functional connectivity in the motor cortex of resting human brain using echo-planar MRI. Magn. Reson. Med. 34, 537–541.

Blondel, V.D., Guillaume, J.-L., Lambiotte, R., Lefebvre, E., 2008. Fast unfolding of communities in large networks. J. Stat. Mech. 2008, P10008.

Boynton, G.M., Engel, S.A., Glover, G.H., Heeger, D.J., 1996. Linear systems analysis of functional magnetic resonance imaging in human V1. J. Neurosci. 16, 4207–4221.

Braga, R.M., Buckner, R.L., 2017. Parallel Interdigitated Distributed Networks within the Individual Estimated by Intrinsic Functional Connectivity. Neuron 95, 457–471.e5.

Broca, P., 1861. Remarques sur le siège de la faculté du langage articulé, suivies d’une observation d’aphémie (perte de la parole). Bulletin et Memoires de la Societe anatomique de Paris 6, 330–357.

Buckner, R.L., Krienen, F.M., Castellanos, A., Diaz, J.C., Yeo, B.T.T., 2011. The organization of the human cerebellum estimated by intrinsic functional connectivity. J. Neurophysiol. 106, 2322–2345.

Bullmore, E., Sporns, O., 2009. Complex brain networks: graph theoretical analysis of structural and functional systems. Nat. Rev. Neurosci. 10, 186–198.

Bürgel, U., Amunts, K., Hoemke, L., Mohlberg, H., Gilsbach, J.M., Zilles, K., 2006. White matter fiber tracts of the human brain: three-dimensional mapping at microscopic resolution, topography and intersubject variability. Neuroimage 29, 1092–1105.

Camara, E., Krämer, U.M., Cunillera, T., Marco-Pallarés, J., Cucurell, D., Nager, W., Mestres-Missé, A., Bauer, P., Schüle, R., Schöls, L., Tempelmann, C., Rodriguez-Fornells, A., Münte, T.F., 2010. The effects of COMT (Val108/158Met) and DRD4 (SNP-521) dopamine genotypes on brain activations related to valence and magnitude of rewards. Cereb. Cortex 20, 1985–1996.

Choi, E.Y., Yeo, B.T.T., Buckner, R.L., 2012. The organization of the human striatum estimated by intrinsic functional connectivity. J. Neurophysiol. 108, 2242–2263.

Clopper, C.J., Pearson, E.S., 1934. The Use of Confidence or Fiducial Limits Illustrated in the Case of the Binomial. Biometrika 26, 404–413.

Cole, M.W., Bassett, D.S., Power, J.D., Braver, T.S., Petersen, S.E., 2014a. Intrinsic and task-evoked network architectures of the human brain. Neuron 83, 238–251.

Cole, M.W., Etzel, J.A., Zacks, J.M., Schneider, W., Braver, T.S., 2011. Rapid transfer of abstract rules to novel contexts in human lateral prefrontal cortex. Front. Hum. Neurosci. 5, 142.

Cole, M.W., Ito, T., Bassett, D.S., Schultz, D.H., 2016a. Activity flow over resting-state networks shapes cognitive task activations. Nat. Neurosci. 19, 1718–1726.

Cole, M.W., Repovš, G., Anticevic, A., 2014b. The frontoparietal control system: a central role in mental health. Neuroscientist 20, 652–664.

Cole, M.W., Reynolds, J.R., Power, J.D., Repovs, G., Anticevic, A., Braver, T.S., 2013. Multi-task connectivity reveals flexible hubs for adaptive task control. Nat. Neurosci. 16, 1348–1355.

Cole, M.W., Yang, G.J., Murray, J.D., Repovš, G., Anticevic, A., 2016b. Functional connectivity change as shared signal dynamics. J. Neurosci. Methods 259, 22–39.

De Baene, W., Vogels, R., 2010. Effects of Adaptation on the Stimulus Selectivity of Macaque Inferior Temporal Spiking Activity and Local Field Potentials. Cereb. Cortex 20, 2145–2165.

Deichmann, R., Gottfried, J.A., Hutton, C., Turner, R., 2003. Optimized EPI for fMRI studies of the orbitofrontal cortex. Neuroimage 19, 430–441.

Delgado, M.R., Nystrom, L.E., Fissell, C., Noll, D.C., Fiez, J.A., 2000. Tracking the hemodynamic responses to reward and punishment in the striatum. J. Neurophysiol. 84, 3072–3077.

Dosenbach, N.U.F., Fair, D.A., Miezin, F.M., Cohen, A.L., Wenger, K.K., Dosenbach, R.A.T., Fox, M.D., Snyder, A.Z., Vincent, J.L., Raichle, M.E., Schlaggar, B.L., Petersen, S.E., 2007. Distinct brain networks for adaptive and stable task control in humans. Proceedings of the National Academy of Sciences 104, 11073–11078.

Doucet, G., Naveau, M., Petit, L., Delcroix, N., Zago, L., Crivello, F., Jobard, G., Tzourio-Mazoyer, N., Mazoyer, B., Mellet, E., Joliot, M., 2011. Brain activity at rest: a multiscale hierarchical functional organization. J. Neurophysiol. 105, 2753–2763.

Duncan, J., 2010. The multiple-demand (MD) system of the primate brain: mental programs for intelligent behaviour. Trends Cogn. Sci. 14, 172–179.

Eichenbaum, H., Yonelinas, A.P., Ranganath, C., 2007. The medial temporal lobe and recognition memory. Annu. Rev. Neurosci. 30, 123–152.

Eickhoff, S.B., Stephan, K.E., Mohlberg, H., Grefkes, C., Fink, G.R., Amunts, K., Zilles, K., 2005. A new SPM toolbox for combining probabilistic cytoarchitectonic maps and functional imaging data. Neuroimage 25, 1325–1335.

Fan, L., Li, H., Zhuo, J., Zhang, Y., Wang, J., Chen, L., Yang, Z., Chu, C., Xie, S., Laird, A.R., Fox, P.T., Eickhoff, S.B., Yu, C., Jiang, T., 2016. The Human Brainnetome Atlas: A New Brain Atlas Based on Connectional Architecture. Cereb. Cortex 26, 3508–3526.

Feinberg, D.A., Moeller, S., Smith, S.M., Auerbach, E., Ramanna, S., Glasser, M.F., Miller, K.L., Ugurbil, K., Yacoub, E., 2010. Multiplexed Echo Planar Imaging for Sub-Second Whole Brain FMRI and Fast Diffusion Imaging. PLoS One 5, e15710.

Felleman, D., Van Essen, D., 1991. Distributed hierarchical processing in the primate cerebral cortex. Cereb. Cortex 1, 1–47.

Fiorillo, C.D., Tobler, P.N., Schultz, W., 2003. Discrete coding of reward probability and uncertainty by dopamine neurons. Science 299, 1898–1902.

Fox, M.D., Snyder, A.Z., Vincent, J.L., Corbetta, M., Essen, D.C.V., Raichle, M.E., 2005. The human brain is intrinsically organized into dynamic, anticorrelated functional networks. Proc. Natl. Acad. Sci. U. S. A. 102, 9673–9678.

Fritsch, G., Hitzig, E., 1870. Electric excitability of the cerebrum (Uber die elektrische Erregbarkeit des Grosshirns). Arch Anat Physiol Wissen 37, 300–332.

Gaiteri, C., Chen, M., Szymanski, B., Kuzmin, K., Xie, J., Lee, C., Blanche, T., Chaibub Neto, E., Huang, S.-C., Grabowski, T., Madhyastha, T., Komashko, V., 2015. Identifying robust communities and multi-community nodes by combining top-down and bottom-up approaches to clustering. Sci. Rep. 5, 16361.

Gazzaniga, M.S., 2005. Forty-five years of split-brain research and still going strong. Nat. Rev. Neurosci. 6, 653–659.

Gazzaniga, M.S., Bogen, J.E., Sperry, R.W., 1962. Some functional effects of sectioning the cerebral commissures in man. Proceedings of the National Academy of Sciences 48, 1765–1769.

Girvan, M., Newman, M.E.J., 2002. Community structure in social and biological networks. Proc. Natl. Acad. Sci. U. S. A. 99, 7821–7826.

Glasser, M.F., Coalson, T.S., Bijsterbosch, J.D., Harrison, S.J., Harms, M.P., Anticevic, A., Van Essen, D.C., Smith, S.M., 2017. Using Temporal ICA to Selectively Remove Global Noise While Preserving Global Signal in Functional MRI Data. bioRxiv 193862.

Glasser, M.F., Coalson, T.S., Robinson, E.C., Hacker, C.D., Harwell, J., Yacoub, E., Ugurbil, K., Andersson, J., Beckmann, C.F., Jenkinson, M., Smith, S.M., Van Essen, D.C., 2016. A multi-modal parcellation of human cerebral cortex. Nature 536, 171–178.

Glasser, M.F., Sotiropoulos, S.N., Wilson, J.A., Coalson, T.S., Fischl, B., Andersson, J.L., Xu, J., Jbabdi, S., Webster, M., Polimeni, J.R., Van Essen, D.C., Jenkinson, M., WU-Minn HCP Consortium, 2013. The minimal preprocessing pipelines for the Human Connectome Project. Neuroimage 80, 105–124.

Gordon, E.M., Laumann, T.O., Adeyemo, B., Huckins, J.F., Kelley, W.M., Petersen, S.E., 2016. Generation and Evaluation of a Cortical Area Parcellation from Resting-State Correlations. Cereb. Cortex 26, 288–303.

Gordon, E.M., Laumann, T.O., Gilmore, A.W., Newbold, D.J., Greene, D.J., Berg, J.J., Ortega, M., Hoyt-Drazen, C., Gratton, C., Sun, H., Hampton, J.M., Coalson, R.S., Nguyen, A.L., McDermott, K.B., Shimony, J.S., Snyder, A.Z., Schlaggar, B.L., Petersen, S.E., Nelson, S.M., Dosenbach, N.U.F., 2017. Precision Functional Mapping of Individual Human Brains. Neuron 95, 791–807.e7.

Guimerà, R., Mossa, S., Turtschi, A., Amaral, L.A.N., 2005. The worldwide air transportation network: Anomalous centrality, community structure, and cities’ global roles. Proc. Natl. Acad. Sci. U. S. A. 102, 7794–7799.

Hampson, M., Peterson, B.S., Skudlarski, P., Gatenby, J.C., Gore, J.C., 2002. Detection of functional connectivity using temporal correlations in MR images. Hum. Brain Mapp. 15, 247–262.

Harmelech, T., Preminger, S., Wertman, E., Malach, R., 2013. The Day-After Effect: Long Term, Hebbian-Like Restructuring of Resting-State fMRI Patterns Induced by a Single Epoch of Cortical Activation. Journal of Neuroscience 33, 9488–9497.

Herlenius, E., Lagercrantz, H., 2004. Development of neurotransmitter systems during critical periods. Exp. Neurol. 190 Suppl 1, S8–21.

Jolkkonen, E., Pitkänen, A., 1998. Intrinsic connections of the rat amygdaloid complex: projections originating in the central nucleus. J. Comp. Neurol. 395, 53–72.

Justin Rossi, P., Peden, C., Castellanos, O., Foote, K.D., Gunduz, A., Okun, M.S., 2017. The human subthalamic nucleus and globus pallidus internus differentially encode reward during action control. Hum. Brain Mapp. 38, 1952–1964.

Kahnt, T., Heinzle, J., Park, S.Q., Haynes, J.-D., 2011. Decoding the formation of reward predictions across learning. J. Neurosci. 31, 14624–14630.

Kelly, R.M., Strick, P.L., 2003. Cerebellar loops with motor cortex and prefrontal cortex of a nonhuman primate. J. Neurosci. 23, 8432–8444.

Krienen, F.M., Yeo, B.T.T., Buckner, R.L., 2014. Reconfigurable task-dependent functional coupling modes cluster around a core functional architecture. Philos. Trans. R. Soc. Lond. B Biol. Sci. 369. https://doi.org/10.1098/rstb.2013.0526

Lancichinetti, A., Radicchi, F., Ramasco, J.J., Fortunato, S., 2011. Finding statistically significant communities in networks. PLoS One 6, e18961.

Laumann, T.O., Gordon, E.M., Adeyemo, B., Snyder, A.Z., Joo, S.J., Chen, M.-Y., Gilmore, A.W., McDermott, K.B., Nelson, S.M., Dosenbach, N.U.F., Schlaggar, B.L., Mumford, J.A., Poldrack, R.A., Petersen, S.E., 2015. Functional System and Areal Organization of a Highly Sampled Individual Human Brain. Neuron 87, 657–670.

Linzenbold, W., Lindig, T., Himmelbach, M., 2011. Functional neuroimaging of the oculomotor brainstem network in humans. Neuroimage 57, 1116–1123.

Mantini, D., Corbetta, M., Romani, G.L., Orban, G.A., Vanduffel, W., 2013. Evolutionarily Novel Functional Networks in the Human Brain? J. Neurosci. 33, 3259–3275.

Marchette, S.A., Vass, L.K., Ryan, J., Epstein, R.A., 2014. Anchoring the neural compass: coding of local spatial reference frames in human medial parietal lobe. Nat. Neurosci. 17, 1598–1606.

Marx, E., Deutschländer, A., Stephan, T., Dieterich, M., Wiesmann, M., Brandt, T., 2004. Eyes open and eyes closed as rest conditions: impact on brain activation patterns. Neuroimage 21, 1818–1824.

McAvoy, M., Mitra, A., Coalson, R.S., Petersen, S.E., Raichle, M.E., 2015. Unmasking Language Lateralization in Human Brain Intrinsic Activity. Cereb. Cortex. https://doi.org/10.1093/cercor/bhv007

Merboldt, K.-D., Finsterbusch, J., Frahm, J., 2000. Reducing Inhomogeneity Artifacts in Functional MRI of Human Brain Activation—Thin Sections vs Gradient Compensation. J. Magn. Reson. 145, 184–191.

Mesulam, M.M., 1998. From sensation to cognition. Brain 121 (Pt 6), 1013–1052.

Middleton, F.A., Strick, P.L., 1994. Anatomical evidence for cerebellar and basal ganglia involvement in higher cognitive function. Science 266, 458–461.

Moeller, S., Yacoub, E., Olman, C.A., Auerbach, E., Strupp, J., Harel, N., Uğurbil, K., 2010. Multiband multislice GE-EPI at 7 tesla, with 16-fold acceleration using partial parallel imaging with application to high spatial and temporal whole-brain fMRI. Magn. Reson. Med. 63, 1144–1153.

Power, J.D., Cohen, A.L., Nelson, S.M., Wig, G.S., Barnes, K.A., Church, J.A., Vogel, A.C., Laumann, T.O., Miezin, F.M., Schlaggar, B.L., Petersen, S.E., 2011. Functional network organization of the human brain. Neuron 72, 665–678.

Power, J.D., Laumann, T.O., Plitt, M., Martin, A., Petersen, S.E., 2017a. On Global fMRI Signals and Simulations. Trends Cogn. Sci. 21, 911–913.

Power, J.D., Mitra, A., Laumann, T.O., Snyder, A.Z., Schlaggar, B.L., Petersen, S.E., 2014. Methods to detect, characterize, and remove motion artifact in resting state fMRI. Neuroimage 84, 320–341.

Power, J.D., Petersen, S.E., 2013. Control-related systems in the human brain. Curr. Opin. Neurobiol. 23, 223–228.

Power, J.D., Plitt, M., Laumann, T.O., Martin, A., 2017b. Sources and implications of whole-brain fMRI signals in humans. Neuroimage 146, 609–625.

Power, J.D., Schlaggar, B.L., Lessov-Schlaggar, C.N., Petersen, S.E., 2013. Evidence for hubs in human functional brain networks. Neuron 79, 798–813.

Robinson, E.C., Jbabdi, S., Glasser, M.F., Andersson, J., Burgess, G.C., Harms, M.P., Smith, S.M., Van Essen, D.C., Jenkinson, M., 2014. MSM: a new flexible framework for Multimodal Surface Matching. Neuroimage 100, 414–426.

Rogers, T.T., Hocking, J., Noppeney, U., Mechelli, A., Gorno-Tempini, M.L., Patterson, K., Price, C.J., 2006. Anterior temporal cortex and semantic memory: reconciling findings from neuropsychology and functional imaging. Cogn. Affect. Behav. Neurosci. 6, 201–213.

Rosvall, M., Bergstrom, C.T., 2008. Maps of random walks on complex networks reveal community structure. Proceedings of the National Academy of Sciences 105, 1118–1123.

Rubinov, M., Sporns, O., 2011. Weight-conserving characterization of complex functional brain networks. Neuroimage 56, 2068–2079.

Rubinov, M., Sporns, O., 2010. Complex network measures of brain connectivity: uses and interpretations. Neuroimage 52, 1059–1069.

Ryali, S., Chen, T., Supekar, K., Menon, V., 2012. Estimation of functional connectivity in fMRI data using stability selection-based sparse partial correlation with elastic net penalty. Neuroimage 59, 3852–3861.

Salimi-Khorshidi, G., Douaud, G., Beckmann, C.F., Glasser, M.F., Griffanti, L., Smith, S.M., 2014. Automatic denoising of functional MRI data: combining independent component analysis and hierarchical fusion of classifiers. Neuroimage 90, 449–468.

Schultz, W., 2006. Behavioral theories and the neurophysiology of reward. Annu. Rev. Psychol. 57, 87–115.

Schultz, W., Apicella, P., Scarnati, E., Ljungberg, T., 1992. Neuronal activity in monkey ventral striatum related to the expectation of reward. J. Neurosci. 12, 4595–4610.

Seeley, W.W., Menon, V., Schatzberg, A.F., Keller, J., Glover, G.H., Kenna, H., Reiss, A.L., Greicius, M.D., 2007. Dissociable intrinsic connectivity networks for salience processing and executive control. J. Neurosci. 27, 2349–2356.

Smith, A.T., Cotton, P.L., Bruno, A., Moutsiana, C., 2009. Dissociating vision and visual attention in the human pulvinar. J. Neurophysiol. 101, 917–925.

Smith, S.M., Beckmann, C.F., Andersson, J., Auerbach, E.J., Bijsterbosch, J., Douaud, G., Duff, E., Feinberg, D.A., Griffanti, L., Harms, M.P., Kelly, M., Laumann, T., Miller, K.L., Moeller, S., Petersen, S., Power, J., Salimi-Khorshidi, G., Snyder, A.Z., Vu, A.T., Woolrich, M.W., Xu, J., Yacoub, E., Uğurbil, K., Van Essen, D.C., Glasser, M.F., WU-Minn HCP Consortium, 2013. Resting-state fMRI in the Human Connectome Project. Neuroimage 80, 144–168.

Smith, S.M., Fox, P.T., Miller, K.L., Glahn, D.C., Fox, P.M., Mackay, C.E., Filippini, N., Watkins, K.E., Toro, R., Laird, A.R., Beckmann, C.F., 2009. Correspondence of the brain’s functional architecture during activation and rest. Proc. Natl. Acad. Sci. U. S. A. 106, 13040–13045.

Sporns, O., 2014. Contributions and challenges for network models in cognitive neuroscience. Nat. Neurosci. 17, 652–660.

Stark, D.E., Margulies, D.S., Shehzad, Z.E., Reiss, P., Kelly, A.M.C., Uddin, L.Q., Gee, D.G., Roy, A.K., Banich, M.T., Castellanos, F.X., Milham, M.P., 2008. Regional variation in interhemispheric coordination of intrinsic hemodynamic fluctuations. J. Neurosci. 28, 13754–13764.

Stokes, M.G., Kusunoki, M., Sigala, N., Nili, H., Gaffan, D., Duncan, J., 2013. Dynamic coding for cognitive control in prefrontal cortex. Neuron 78, 364–375.

Traud, A., Kelsic, E., Mucha, P., Porter, M., 2011. Comparing Community Structure to Characteristics in Online Collegiate Social Networks. SIAM Rev. 53, 526–543.

Uğurbil, K., Xu, J., Auerbach, E.J., Moeller, S., Vu, A.T., Duarte-Carvajalino, J.M., Lenglet, C., Wu, X., Schmitter, S., Van de Moortele, P.F., Strupp, J., Sapiro, G., De Martino, F., Wang, D., Harel, N., Garwood, M., Chen, L., Feinberg, D.A., Smith, S.M., Miller, K.L., Sotiropoulos, S.N., Jbabdi, S., Andersson, J.L.R., Behrens, T.E.J., Glasser, M.F., Van Essen, D.C., Yacoub, E., 2013. Pushing spatial and temporal resolution for functional and diffusion MRI in the Human Connectome Project. Neuroimage, Mapping the Connectome 80, 80–104.

van Baarsen, K.M., Kleinnijenhuis, M., Jbabdi, S., Sotiropoulos, S.N., Grotenhuis, J.A., van Cappellen van Walsum, A.M., 2016. A probabilistic atlas of the cerebellar white matter. Neuroimage 124, 724–732.

Van Essen, D.C., Glasser, M.F., 2014. In vivo architectonics: a cortico-centric perspective. Neuroimage 93 Pt 2, 157–164.

Van Essen, D.C., Smith, S.M., Barch, D.M., Behrens, T.E.J., Yacoub, E., Ugurbil, K., 2013. The WU-Minn Human Connectome Project: An overview. Neuroimage, Mapping the Connectome 80, 62–79.

Wallis, J.D., Miller, E.K., 2003. Neuronal activity in primate dorsolateral and orbital prefrontal cortex during performance of a reward preference task. Eur. J. Neurosci. 18, 2069–2081.

Wang, D., Buckner, R.L., Fox, M.D., Holt, D.J., Holmes, A.J., Stoecklein, S., Langs, G., Pan, R., Qian, T., Li, K., Baker, J.T., Stufflebeam, S.M., Wang, K., Wang, X., Hong, B., Liu, H., 2015. Parcellating cortical functional networks in individuals. Nat. Neurosci. 18, 1853–1860.

Wernicke, C., 1874. Der aphasische Symptomencomplex; eine psychologische Studie auf anatomischer Basis. Cohn & Weigert, Breslau.

Yeo, B.T.T., Krienen, F.M., Eickhoff, S.B., Yaakub, S.N., Fox, P.T., Buckner, R.L., Asplund, C.L., Chee, M.W.L., 2015. Functional Specialization and Flexibility in Human Association Cortex. Cereb. Cortex 25, 3654–3672.

Yeo, B.T.T., Krienen, F.M., Sepulcre, J., Sabuncu, M.R., Lashkari, D., Hollinshead, M., Roffman, J.L., Smoller, J.W., Zöllei, L., Polimeni, J.R., Fischl, B., Liu, H., Buckner, R.L., 2011. The organization of the human cerebral cortex estimated by intrinsic functional connectivity. J. Neurophysiol. 106, 1125–1165.

Zhang, D., Snyder, A.Z., Fox, M.D., Sansbury, M.W., Shimony, J.S., Raichle, M.E., 2008. Intrinsic functional relations between human cerebral cortex and thalamus. J. Neurophysiol. 100, 1740–1748.

